# Prolonged and distributed processing of facial identity in the human brain

**DOI:** 10.1101/2021.06.23.449599

**Authors:** Rico Stecher, Ilkka Muukkonen, Viljami Salmela, Sophie-Marie Rostalski, Géza Gergely Ambrus, Gyula Kovács

**Affiliations:** Institute of Psychology, Friedrich Schiller University Jena, Jena, Germany; Department of Psychology and Logopedics, University of Helsinki, Finland

**Keywords:** face identification, representational similarity analysis, EEG-fMRI fusion

## Abstract

The recognition of facial identity is essential for social interactions. Despite extensive prior fMRI and EEG/MEG research on the neural representations of familiar faces, we know little about the spatio-temporal dynamics of face identity information. Therefore, we applied a novel multimodal approach by fusing the neuronal responses recorded in an fMRI and an EEG experiment. We analyzed the neural responses to naturally varying famous faces and traced how face identity emerges over time in different areas of the brain. We found that image invariant face identity information prevails over an extended time period (from 150 to 810 ms after stimulus onset) in the representational geometry of a broadly distributed network of parietal, temporal, and frontal areas with overlapping temporal profiles. These results challenge the current hierarchical models of face perception and suggest instead concerted and parallel activation of multiple nodes in the brain’s identity coding network while processing information of familiar faces.

## Introduction

Through the development of multivariate pattern analysis (MVPA) techniques, we have gained a deeper insight in the underlying neural mechanisms of face recognition and the development of face familiarity. Previous neuroimaging studies, focusing on the spatial domain, have shown that information, which is relevant for the identification of familiar faces, can be decoded from various regions of the face processing network (Gobbini and Haxby, 2007). These areas include parts of the core-network, processing various perceptual face aspects, such as the fusiform face area (FFA; Anzellotti et al., 2014; Axelrod and Yovel, 2015; Gilaie-Dotan and Malach, 2007; Goesaert and Op de Beeck, 2013; Nestor et al., 2011; Verosky et al., 2013; Weibert et al., 2016), the anterior temporal lobe (ATL; Anzellotti et al., 2014; Kriegeskorte et al., 2007; Nasr & Tootell, 2012), and also parts of the so-called extended network, such as the medial temporal regions, medial and lateral parietal regions, the amygdala, the insula, and medial and inferior frontal regions (for a recent review see Kovács, 2020). On the other hand, a handful of EEG/MEG studies have used MVPA to investigate the temporal emergence of face identity representations. Identity can be decoded for both unfamiliar (Nemrodov et al., 2018, 2016; Vida et al., 2017) and familiar faces (Ambrus et al., 2019; Dobs et al., 2019) within the first 200 ms post-stimulus onset. Furthermore, there seems to be a difference between the early and the later decoding phases in the sensitivity to low-level image-specific variations of the faces. While the earlier representations of face identity can be explained by broader visual categories that are shared between stimuli, such as sex or age, the representations detected from 400 ms onward are the result of higher-level, image invariant encoding of identity (Ambrus et al., 2019; Wiese et al., 2019).

Although a large body of research indicates that the several-hundred millisecond long activation of an extensive cortical network is necessary for the correct recognition of face identity, the exact temporal dynamics of the network is still largely unexplored. Dobs et al. (2019), using familiar and unfamiliar faces differing in age and sex, showed that basic visual dimensions of a face, such as low-level image-specific features but also information like age, sex and identity are decodable from MEG signals at different processing stages. These results suggest a coarse-to-fine information processing trajectory. Thus, it is possible that the higher-level information within a familiar face (such as their image-independent perceptual representations, the recall of person-related semantic and episodic memories, the encoding of personality traits and attitudes as well as the associated emotional responses) are also extracted at different processing stages. Alternatively, the activation of the various areas underlying the above dimensions could occur in a temporally overlapping manner, parallel to each other.

To reveal the spatio-temporal dynamics of face identity processing, we used representational similarity analysis (RSA; Kriegeskorte et al, 2017; Kriegeskorte and Kievit, 2013) to integrate fMRI and EEG data. By relating multivariate similarity spaces obtained from EEG and fMRI via RSA, we were able to combine the superior temporal resolution of EEG with the precise spatial resolution of fMRI. This cutting-edge analysis method, that has only been used in a hand-full of studies so far, has brought significant advances in understanding the processing of objects (Cichy et al., 2016, 2014; Cichy and Pantazis, 2017; Hebart et al., 2018), biological motion (Chang et al., 2021) as well as emotions present in voices (Giordano et al., 2021) or faces (Muukkonen et al.,2020; Bayer et al, 2021).

Here we used this approach to investigate the encoding of ambient faces (Jenkins et al., 2011) of famous persons across space and time. Our results show distributed and prolonged processing of face identity, starting around 150 ms after stimulus onset within the inferior parietal cortex and quickly spreading simultaneously across the entire extended face network.

## Methods

### Participants

24 healthy participants took part in the study. The data of four participants was excluded from the final analysis due to excessive head movements during the fMRI measurement, resulting in a poor signal-to-noise ratio in the individual data sets. Thus, the data of 20 participants (10 male) with a mean age of 22.05 years (SD = 2.26) was analyzed. This sample size was based on previous studies employing the representational-similarity-based fusion of M/EEG and fMRI (Cichy et al., 2014; Muukkonen et al., 2020), but was slightly increased to enhance statistical power. All participants had normal or corrected-to-normal eyesight. Participants were either compensated financially or with partial course credits. The study was conducted in accordance with the Declaration of Helsinki and was accepted by the ethics committee of the Friedrich Schiller University Jena.

### EEG task and procedure

The participants’ task was identical to the one employed by Ambrus et al. (2019). Briefly, participants viewed highly varying, ambient color images of four celebrities on a uniform gray backround (Angelina Jolie, Heidi Klum, Leonardo DiCaprio and Til Schweiger; size: 4.4° visual angle). Every participant could correctly identify these celebrities by recalling their names and occupations. During the experiment, participants were presented with 10 images per identity (4 x 10 = 40 in total) in random order. Each stimulus was repeated 40 times (1600 non-target trials in total). In order to maintain the participants’ attention, 160 target trials were also included in which the images were rotated clockwise or counterclockwise (10°) and participants were required to signal the appearance of these images by pressing a button. These target trials were excluded from any further analysis. The experiment was written in Psychopy (Peirce, 2008).

### EEG data acquisition and pre-processing

The EEG was recorded in a dimly lit, electrically shielded chamber using a 64-channel BioSemi Active II system (BioSemi, Amsterdam, Netherlands) with a sample rate of 512 Hz. An electrooculogram (EOG) was recorded from the outer canthi of both eyes as well as above and below the left eye. Participants were required to place their heads on a chin rest to ensure a distance of 96 cm between their eyes and the computer screen.

The EEG data was pre-processed in EEGLAB (Delorme and Makeig, 2004). First, data was re-referenced to common average, then a band-pass filter between 0.1 and 70 Hz was applied, and line noise was removed with the CleanLine plugin (Mullen, 2012). No further artifact correction or rejection was performed, as classifiers are thought to be robust to noise in time-series neuroimaging data (Grootswagers et al., 2017). Data was downsampled to 100 Hz to increase signal-to-noise ratio. Finally, the EEG data was segmented between -200 ms and 1300 ms relative to stimulus onset and then baseline corrected.

### fMRI task and procedure

We always performed the fMRI recording before the EEG recording session and participants were only invited to the EEG session if they did not show excessive head movements during the fMRI measurements.

A modified version of the EEG paradigm was used during the fMRI recording sessions. Participants completed four functional runs: In each run, every image was presented 5 times resulting in 800 non-target trials in total. During each trial, the images (size: 3.4° visual angle) were presented for 600 ms, followed by an interstimulus interval (ISI) of either 1400 or 3400 ms (with equal probability) during which a black fixation cross was shown. Thus, the total trial length was either 2000 ms or 4000 ms. The position of the images was jittered by 10 pixels along the x- and y-axes across trials to avoid low-level adaptation. We included 4 target trials per run (16 in total) in which the participants had to respond to images that were rotated clockwise by 10° by pressing a button. All target trials were modelled in SPM as a nuisance regressor, but again discarded from any further analysis. The experiment was programmed in MATLAB (Mathworks, 2013), using PsychToolbox (Brainard, 1997; Kleiner et al., 2007; Pelli, 1997).

### fMRI data acquisition and pre-processing

fMRI data was recorded using a Prisma fit 3T scanner (Siemens Healthcare, Erlangen, Germany), fitted with a 20-channel head coil. High-resolution T1-weighted anatomical scans were collected with an MP-RAGE sequence (TR = 2300 ms, TE = 3.03 ms, 192 slices, 1 mm isotropic voxel size). Functional scans were obtained using a T2*-weighted EPI sequence (35 slices, 10° tilted relative to axial, TR = 2000 ms, echo time (TE) = 30 ms, flip angle 90°, in plane resolution 3 mm isotopic voxel size). Our analysis pipeline consisted of slice-timing, realignment, co-registration to the anatomical scan, normalization to MNI-152 space and resampling to 2 x 2 x 2 mm resolution. We pre-processed the fMRI data in SPM12 (www.fil.ion.ucl.ac.uk/spm/).

### Representational-similarity-based EEG-fMRI fusion

In order to trace identity information across both space and time, we conducted a representational-similarity-based EEG-fMRI fusion (Figure 1; Cichy et al., 2014; Cichy and Pantazis, 2017; Muukkonen et al., 2020). First, we created neural representational dissimilarity matrices (RDMs) for each time point of the segmented EEG data and every voxel in the functional MRI images.

**Figure 1.**
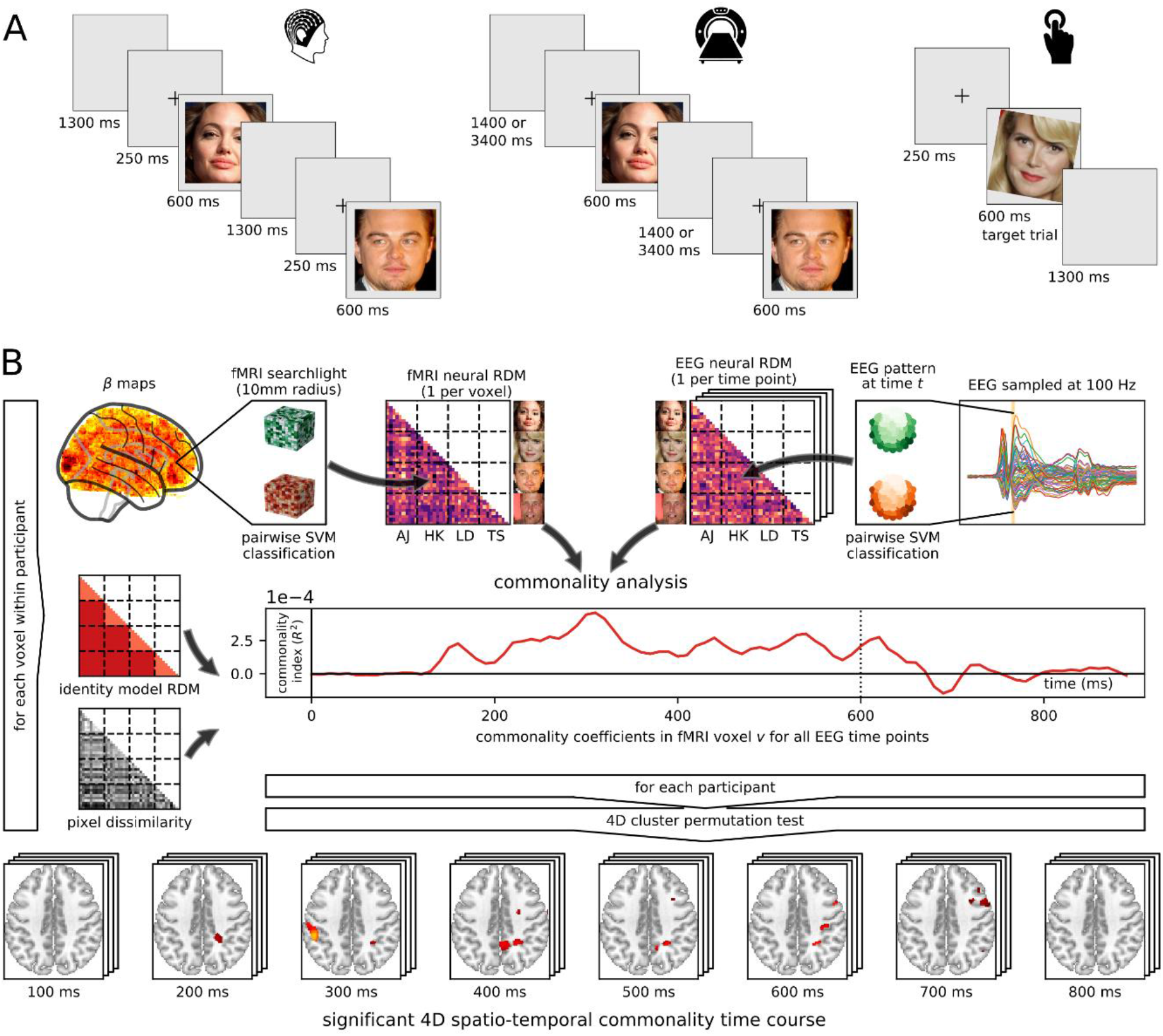
Design and analysis approach. A. Spatial (fMRI) and temporal (EEG) RDMs were constructed respectively for each voxel (10 mm searchlight radius) and each time point (10 ms). Each cell of these RDMs reflects the pairwise decoding accuracy between a given stimulus pair, obtained from the corresponding neural data for each of the participants. B. Using commonality analysis, the unique variance shared between the participant-specific fMRI RDM at a given voxel, the averaged EEG RDM at a given time point and the identity model RDM was determined, while controlling for the pixel dissimilarity RDM. C. This yielded a 4D spatio-temporal commonality time course for each participant, revealing the spatio-temporal dynamics of face identity representations.

We constructed the EEG RDMs by conducting single-trial pairwise decoding of every possible stimulus pair with 5-fold-cross-validation, separately for each participant. We assigned 20% of the trials of a given stimulus to 1 of 5 datasets randomly, so that each set contained an even number of trials per stimulus. Support vector machine (SVM) classifiers were then trained on 4 of the datasets and tested on the 5th in a leave-one-out design. This was done separately at each time point and for each possible stimulus pair with all of the 64 EEG channels as features. The classification accuracy at each time point was then calculated as the average of all cross-validation folds. We assembled the resulting pairwise decoding accuracies for each possible stimulus pair into 40 x 40 RDMs, one for each time point, omitting the diagonal to avoid the artificial inflation of similarity between RDMs (Ritchie et al., 2017). To increase the signal-to-noise ratio, we averaged these EEG RDMs at each time point across participants and smoothed the pairwise decoding accuracies with a 30 ms rolling average (Cichy et al., 2016; Muukkonen et al., 2020). All multivariate pattern classifications on the EEG data set were conducted using the Decision Decoding Toolbox (DDTBOX; Bode et al., 2019).

For the classification, L2-norm SVMs with a regularization parameter of C = 1 were used, and were implemented in LIBSVM (Chang and Lin, 2011).

To obtain the fMRI RDMs, we first estimated general linear models (GLMs) for each participant separately, specifying each of the 40 stimuli as a regressor (also including the target trials as a nuisance regressor). The resulting beta maps were then used to create RDMs at each voxel for each participant. We examined each voxel using a searchlight approach (10 mm radius) and trained SVM classifiers to distinguish the activation patterns of each possible combination of stimulus pairs (Kriegeskorte et al., 2006, 2007). We employed a leave-one-run-out design, so in each cross-validation fold, the classifiers were trained on the data of 3 runs and tested on the data of the 4th. The pairwise decoding accuracies were then calculated as the averages across all cross-validation folds. Here again, the decoding accuracies of every possible stimulus pair were combined into 40 x 40 RDMs. All fMRI multivariate pattern classifications were carried out using The Decoding Toolbox (Hebart et al., 2015). Equal to the analysis of the EEG data, we used L2-norm SVMs as classifiers, with a regularization parameter of C = 1, which were implemented in LIBSVM (Chang and Lin, 2011).

Finally, a model-based EEG-fMRI fusion using commonality analysis was conducted (Hebart et al., 2018; Seibold and McPhee, 1979). Commonality analysis is a variance decomposition method in which so-called commonality coefficients are calculated. These are determination coefficients (R^2^) that reflect how much variance is uniquely shared between multiple variables. In this particular application, they allow us to quantify how much variance is uniquely shared among the EEG RDMs at each time point, the fMRI RDMs at each voxel and the model RDMs. We created a conceptual identity model RDM (Figure 1) in which all images of the same identity were considered similar (represented by a 0) and all images of different identities were considered dissimilar (represented by a 1).

In order to control for low-level image similarity, we also created a pixel dissimilarity RDM in the dissimilarity between pixel patterns of each possible stimulus pair was quantified, using 1 - correlation as the distance measure. The approach was identical to Ambrus et al. (2019). Next, we conducted the fusion analysis by first vectorizing the lower triangle of each EEG RDM and fMRI RDM for each participant as well as the identity model RDM and the pixel dissimilarity RDM. We then iterated through the EEG time series and computed the commonality coefficients between the averaged EEG RDM at that specific point in time, the participant-specific fMRI RDM at each voxel and the identity model RDM while controlling for pixel dissimilarity. For each time point-voxel pair we first calculated two squared semipartial Spearman correlation coefficients (R^2^) between the EEG and fMRI RDMs, one in which only the pixel dissimilarity RDM was controlled for and one in which both the identity model RDM and pixel dissimilarity RDM were controlled for. We then obtained the commonality coefficients by subtracting the latter R^2^ from the former. These commonality coefficients thus reflect the variance shared between the EEG, the fMRI RDMs and the identity model RDM, when controlling for pixel dissimilarity.

Note that when calculating these correlations, the EEG RDMs were always the dependent variable. Repeating this procedure for each time-point-voxel-combination and participant resulted in subject-specific 4D spatio-temporal commonality time courses. These consisted of a 3D commonality map for every time point, which contained the spatio-temporal dynamics of face identity information processing. Since we averaged the EEG RDMs at each time point across participants, all between-participant variance in these time courses can solely be attributed to the variance in the participant-specific fMRI RDMs. All 3D commonality maps were smoothed with an 8 mm FWHM Gaussian kernel in SPM12.

### Statistical inference

The obtained 4D spatio-temporal commonality time courses were tested for significance using a non-parametric 4D cluster-permutation test (Nichols and Holmes, 2002). The cluster definition threshold was determined by first conducting a one-sample t-test on each voxel in the 4D spatio-temporal commonality time courses in the baseline period (-200 to 0 ms). The resulting t-values were then aggregated into one empirical distribution under the null hypothesis and the cluster definition threshold was defined as the 99.5th percentile of this distribution (equivalent to a significance level of p = 0.005; Cichy et al., 2016). For the purpose of constructing a permutation distribution of maximum spatio-temporal cluster sizes, we randomly flipped the sign of each participant’s 4D spatio-temporal commonality time course 5000 times (including the original permutation). During each permutation, a t-map of the 4D spatio-temporal commonality time courses was constructed. This t-map was limited at the cluster definition threshold and 4D spatio-temporal clusters were formed. These were defined as above-threshold voxels that were either spatially and/or temporally contiguous. Finally, the maximum cluster size was determined during each permutation. This resulted in a permutation distribution of maximum cluster sizes under the null hypothesis. The p-value of each spatio-temporal cluster was defined as the proportion of maximum spatio-temporal cluster sizes in the permutation distribution that were at least as large as the cluster itself (Nichols and Holmes, 2002). Spatio-temporal clusters with p < 0.05 were reported as significant. For the sake of concision and interpretability, we applied a spatial and temporal cluster extent threshold to the spatial clusters within the significant spatio-temporal clusters. We only report spatial clusters within the above-defined spatio-temporal clusters that 1) were larger than the median cluster size of all spatial clusters in the spatio-temporal cluster and 2) lasted for at least 30 ms (a spatial cluster was defined as continuous for 30 ms, if at least 10% of its voxels were present at the previous time point and at least 10% were present at the subsequent time point).

The data of the original significant spatio-temporal time course are presented in Figure S1 in the Supplementary Material where the link to an animation of this time course can also be found.

The spatial clusters at each time point within the significant spatio-temporal clusters were at first automatically labeled using the probabilistic cytoarchitectonic maps available in the SPM Anatomy toolbox (Eickhoff et al., 2005). They were then manually double-checked, based on the literature and publicly available brain atlases.

## Results

The fusion of EEG and fMRI data revealed both where and when in the brain the neural activity is associated with facial identity processing. The Supplementary Material contains a table (Table S2) that lists the coordinates and anatomical labels of all areas in the significant spatio-temporal time course. Furthermore, it also contains visualizations (Figure S3) and animations of the areas showing identity-model-explained representational correspondence of EEG and fMRI with a 10 ms temporal resolution. Figure 2b shows schematically, that the earliest face identity representations occur at around 150 ms. Surprisingly, the identity representations in this early time period of the EEG signal correlated with the activity patterns of the right inferior parietal lobule (IPL; Figure 2a). This correlation remained significant from 150 ms to 280 ms, peaked later on at around 550 ms, and then with brief interruptions persisted until about 750 ms.

**Figure 2.**
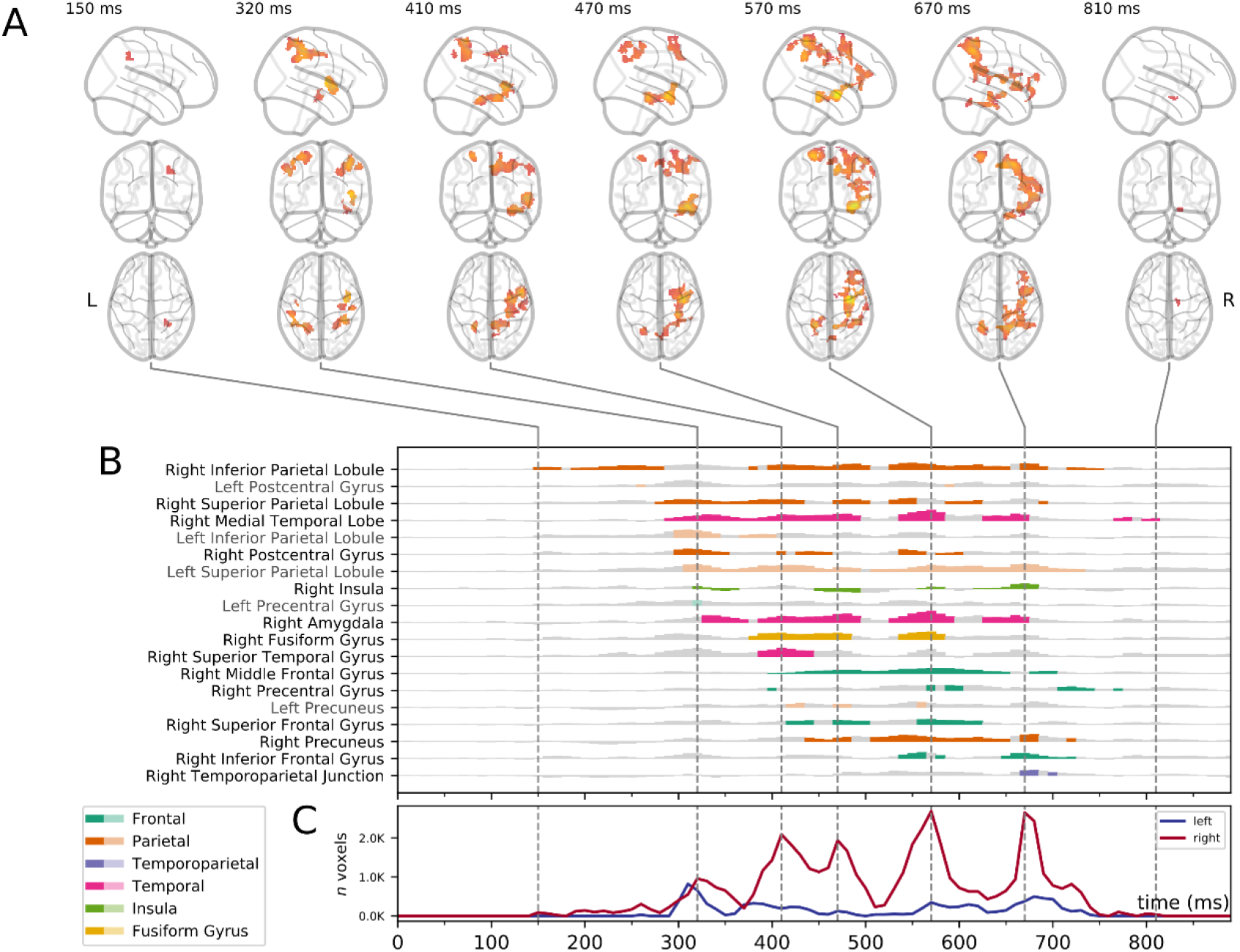
Spatio-temporal RSA of the representations of face identity. A. Significant spatio-temporal identity representations at selected time points. Maps are thresholded for significance (4D cluster-permutation test, n = 20, cluster definition threshold p < 0.005, cluster threshold p < 0.05) and plotted on the MNI template brain. Color scale signals unique variance shared between the fMRI RDM at a given voxel, the EEG RDM at a given time point and the identity model RDM while controlling for pixel dissimilarity. B. Schematic depiction of the identity commonality coefficients for the main regions within the significant spatio-temporal clusters as a function of time (see Table S2 in the Supplementary Material for a detailed list of all areas across time). Time courses reflect the mean identity commonality coefficients within a sphere (10 mm radius) in a given region over time. Colored plots mark time bins in which this region was part of the significant spatio-temporal cluster (see Methods section). C. The sum of voxels within the significant spatio-temporal time course, separately for the right (orange) and left (blue) hemisphere, as a function of time.

Next, starting at around 260-330 ms, significant identity information could be found in different brain regions, including the bilateral superior parietal lobe, left IPL, right medial temporal lobe (MTL), the amygdala, bilateral postcentral gyrus, and the insula. Except for the left IPL (limited between 300 and 400 ms), commonality coefficients of face identity representations in these regions remained periodically significant until 590 to 810 ms after stimulus onset, depending on the respective brain area.

A third wave of activity patterns was detected in cortical areas, including the middle and anterior parts of the right fusiform gyrus as well as the ventral surface of the anterior temporal lobe and the superior temporal gyrus (STG), the right middle and superior frontal areas, the right precentral gyrus and the bilateral precuneus/posterior cingulate (PC/PCC). In these regions, significant commonality coefficients were found, starting from 380 to 440 ms. Face identity information in these areas remained significant at intervals until around 560 to 770 ms. Finally, the right inferior frontal gyrus showed identity-model-explained correspondence between EEG and fMRI data at 540-580 ms and 650-720 ms, while the right temporo-parietal junction (TPJ)/posterior superior temporal sulcus (pSTS) displayed these at 670-700 ms.

Overall, the spatio-temporal structure of face identity encoding showed a strong right-hemispheric lateralization (Figure 2c) activating an extensive network of different areas, with relatively late and extremely long-lasting processing.

It is theoretically possible that the specific activation of some brain regions does not automatically lead to distinct scalp-level EEG patterns and therefore is overlooked by the current fusion analysis. To overcome this limitation we also analysed our fMRI data separately before linking it to EEG. This analysis, however, did not lead to any additional significant clusters encoding identity (RSA with asearchlight with 10 mm radius; 5000 permutations).

## Discussion

The current study aimed at providing detailed information about the spatio-temporal dynamics of face identity processing in the human brain. For this purpose, we integrated fMRI and EEG measurements by using representational similarity analysis (Cichy et al., 2016, 2014). We investigated the involvement of a large variety of brain areas, using a hypothesis-free, searchlight-based decoding approach, rather than limiting the analysis to pre-selected regions deemed relevant in previous works (Kriegeskorte et al., 2007). This analysis enabled us to track the identity-related information flow in the whole cortical face network by combining the temporal precision of EEG with the spatial precision of fMRI. So far, this is the first study to integrate EEG/MEG and fMRI imaging via RSA to study the spatio-temporal dynamics of famous face identity processing. Overall, our analyses show that identity information is present simultaneously in a large network of cortical areas for a long time span, challenging the purely hierarchy-based models of face identity processing.

The first correspondence of fMRI- and EEG-data-derived identity information was found at 150 ms post-stimulus onset in the right IPL. This representation correspondence persisted, with some interruptions, until 750 ms. Such early-onset representations have previously been found in recent EEG/MEG studies for face identity (Dima et al., 2018; Nemrodov et al., 2018, 2016; Vida et al., 2017) as well as for facial expressions (Muukkonen et al., 2020). Our study is the first one to report that image invariant aspects of face identity are initially detectable in the right parietal cortex. Nevertheless, which identity-related features exactly are processed in this area, needs to be explored through systematic investigations in the future.

The spatio-temporal cluster around the rIPL involved the superior intraparietal sulcus (sIPS) as well as the supramarginal and the angular gyri. This region is part of the extended faceprocessing system and encodes gaze-direction and head orientation (for a review, see Haxby et al., 2000) as well as personality traits, attitudes, and mental states of others (Frith and Frith, 1999) and visual short-term memory for objects (Xu and Chun, 2006).

Jeong and Xu (2016) tested the role of the sIPS in object and face identity representation by using MVPA. As stimuli, they used famous faces, varying in viewpoint, hairstyle, expression and age and found significant identity representations in the sIPS around coordinates (MNI [x,y,z,]: 23, -52, 45; Xu and Chun, 2006) which correspond closely to the center of the rIPL cluster of our current study (MNI [x,y,z,]: 34, -48, 36). Therefore, we hypothesize, that the decoding within sIPS is most likely related to abstract identity processing. The persistence of the representational correspondence of the rIPL supports this conclusion, giving rise to the question which role this (previously relatively neglected) parietal area plays in the perception and processing of faces.

Quite surprisingly, in several areas identity-specific representational correspondence arose simultaneously at around 300 ms post-stimulus onset. These included areas that were previously identified as parts of the extended face-processing network (Haxby et al., 2000) and have been shown to be involved in various steps of face identity processing (for a recent review, see Kovács, 2020). First, the information in the rIPL extended bilaterally towards the superior parietal lobule and the postcentral gyrus. These regions were found in a recent meta-analysis to respond to newly learned familiar faces but not to famous or personally familiar faces (Blank et al., 2014). Second, regions of the MTL, including the hippocampus, perirhinal and entorhinal cortex have previously been reported to be involved in storing face-related biographical information and familiarity (Ramon et al., 2015). Third, the amygdala (Ramon et al., 2015) and the insula (Gobbini and Haxby, 2007; Ida Gobbini et al., 2004; Natu and O’Toole, 2011) are related to the enhanced emotional processing of familiar identities. Finally, the sensory function of the precentral gyrus and its contribution to the processing of face identity remains unclear, although studies in the past have shown that it is indeed involved in face perception (Gobbini and Haxby, 2006; Watson et al., 2016).

So far, only a handful of studies used MVPA of fMRI data to locate the identity representations of famous faces. The results of these studies demonstrated the decodability of face identity information in the right FFA (Axelrod and Yovel, 2015) or in a set of occipito-temporal (occipital face area, pSTS) and medial parietal areas, such as the PC/PCC (Tsantani et al., 2019). Accordingly, many of these regions showed identity-specific representational correspondence when combining the EEG and fMRI data. This was also the case for other areas belonging to the core face-processing system (for a different interpretation see Kovács, 2020), namely the ventral surface of the anterior temporal lobe and the superior temporal gyrus (Collins and Olson, 2014; Rajimehr et al., 2009). However, a surprising outcome of the current study is that the representation of identity emerged relatively late, at around 400 ms post-stimulus onset. Recent MVPA studies suggest that the FFA and the ventral anterior temporal areas play a role in the image invariant encoding of face identities (Collins et al., 2016; Goesaert and Op de Beeck, 2013; Nestor et al., 2011; Tsantani et al., 2021). The fact that these areas only showed representational correspondence at a later stage, correspond to their sensitivity to intermediate (Visconti Di Oleggio Castello et al., 2017) or high-level information (Tsantani et al., 2021), which is presumably related to the merging of perceptual and conceptual knowledge about the face (Collins et al., 2016; Morton et al., 2021).

A few frontal and parietal areas showed representational correspondence with an onset of around 400 ms as well: The middle and inferior frontal representational correspondence during this time window might reflect activations of the inferior frontal face area (Axelrod and Yovel, 2013; Chan and Downing, 2011; Collins and Olson, 2014), which is also associated with the view-independent encoding of familiar face identities (Guntupalli et al., 2017). The PC/PCC and other medial parietal regions have also been associated with face familiarity in the past (Gobbini and Haxby, 2006; Visconti Di Oleggio Castello et al., 2017) as well as with the retrieval of person-specific (biographical) information from long-term memory (Burgess et al., 2001; Gilmore et al., 2015; Leech and Sharp, 2014; Wagner et al., 2005).

Lastly, the right pSTS around the TPJ showed representational correspondence between 670-700 ms post-stimulus onset. A recent study, measuring fMRI activation patterns for famous faces and voices found that this region integrates identity representations across visual and acoustic modalities. Therefore, it may serve as a high-level identity processing area (Tsantani et al., 2019), which is consistent with the late occurrence of this correspondence in our analysis.

It is noteworthy that all the above-described identity-selective representations occurred in a strongly right-lateralized network (Figure 2c). This result is in accordance with a wide range of studies suggesting the right-hemisphere dominance of face perception (for a review, see Duchaine and Yovel, 2015). In summary, the above mentioned areas are part of a large network recently defined as the „person identification network” (PIN; Kovács, 2020), which is assumed to be active for famous and, above all, personally familiar faces.

Overall, the spatial pattern suggest the parallel and temporally overlapping activation of several members of the PIN, rather than a coarse-to-fine information processing trajectory. Moreover, the results are congruent with previous studies, proposing the prolonged activation of several processing areas simultaneously for objects (Cichy et al., 2016), emotional voices (Giordano et al., 2021) and facial expressions (Muukkonen et al., 2020). The current study extends these observations to face identity representations.

Importantly, the temporal dynamics of identity representations do not follow the currently accepted hierarchy of processing stages. The identity representation found in areas such as the IPL, which is related to personality, traits and attitudes, preceded that of other areas, suggesting the rapid extraction of high-level information from faces and emphasizing their importance when meeting others. Memory- (medial temporal lobe) and emotion- (amygdala and insula) related representations follow. From around 300 ms and only at around 400 ms identity information in the core network areas can be observed (likely representing higher-level sensory and semantic identity processing via recurrent activations of the network).

It is worth noting that the time window from around 300-400 ms to 600 ms reflects the most robust familiarity-related signals both in univariate (Wiese et al., 2019) and in multivariate studies (Ambrus et al., 2021, 2019; Dobs et al., 2019). This is the time window during which the largest amount of voxels show identity-model-explained representational correspondence with the EEG in our study as well. This emphasizes the importance of the cognitive processes that occur at this stage (presumably memory and emotion processing) for face identity representation. Such an interpretation is in line with current models of PIN (Kovács, 2020) which highlight the role of these regions in separating the processing of familiar and unfamiliar faces.

One open question regards the face specificity of the above pattern of results. This issue is hard to study as it is difficult to find any non-face objects that are as well-known as the faces of our friends and relatives. One such stimulus category may be houses and buildings (Gorno-Tempini and Price, 2001). Indeed, a very recent study compared semantic knowledge for people and places and found separate mechanisms in anterior-temporal and posterior-medial temporal regions but also a joint representation of sematic similarity in the hippocampus (Morton et al, 2021). Testing the identity processing of famous and personally familiar buildings and places might expand the current results over stimulus categories in the future.

In sum, image invariant identity models explain the neural responses to faces in a broadly distributed network. However, our results also show how the representations in the different processing areas develop across time, suggesting prolonged and simultaneous activations of an overlapping network, informing the future development of models of face identification.

## Acknowledgements

The authors would like to thank Louisa Fortwengel, Morgana Dalla Palma, Alexia Dalski, Johannes Lehnen, and Bettina Kamchen for their help in participant recruitment and data acquisition. In addition, they would like to thank Daniel Kaiser for comments on the manuscript and Martin Hebart for advice regarding the analysis. This work was supported by grants from the Deutsche Forschungsgemeinschaft (KO3918/5-1) and from the German Academic Exchange Service (57524936) as well as by a scholarship to R.S. from the Honours Programme of the University of Jena.

## Author Contributions

G.A. and G.K. designed the experiments. R.S. and S.M.R. collected data. G.A., I.M, V.S, R.S. and G.K. designed the analysis. R.S. analyzed the data. G.K., G.A., R.S., I.M., V.S. and S.M.R. wrote the article.

## Data availability

Stimuli and data are available from the Open Science Framework (https://osf.io/8nc3u/). Analysis scripts will be made available upon request.

## Supplementary Material

A) Identity-model-based EEG-fMRI fusion results without spatial and temporal extent threshold

**Figure S1.**
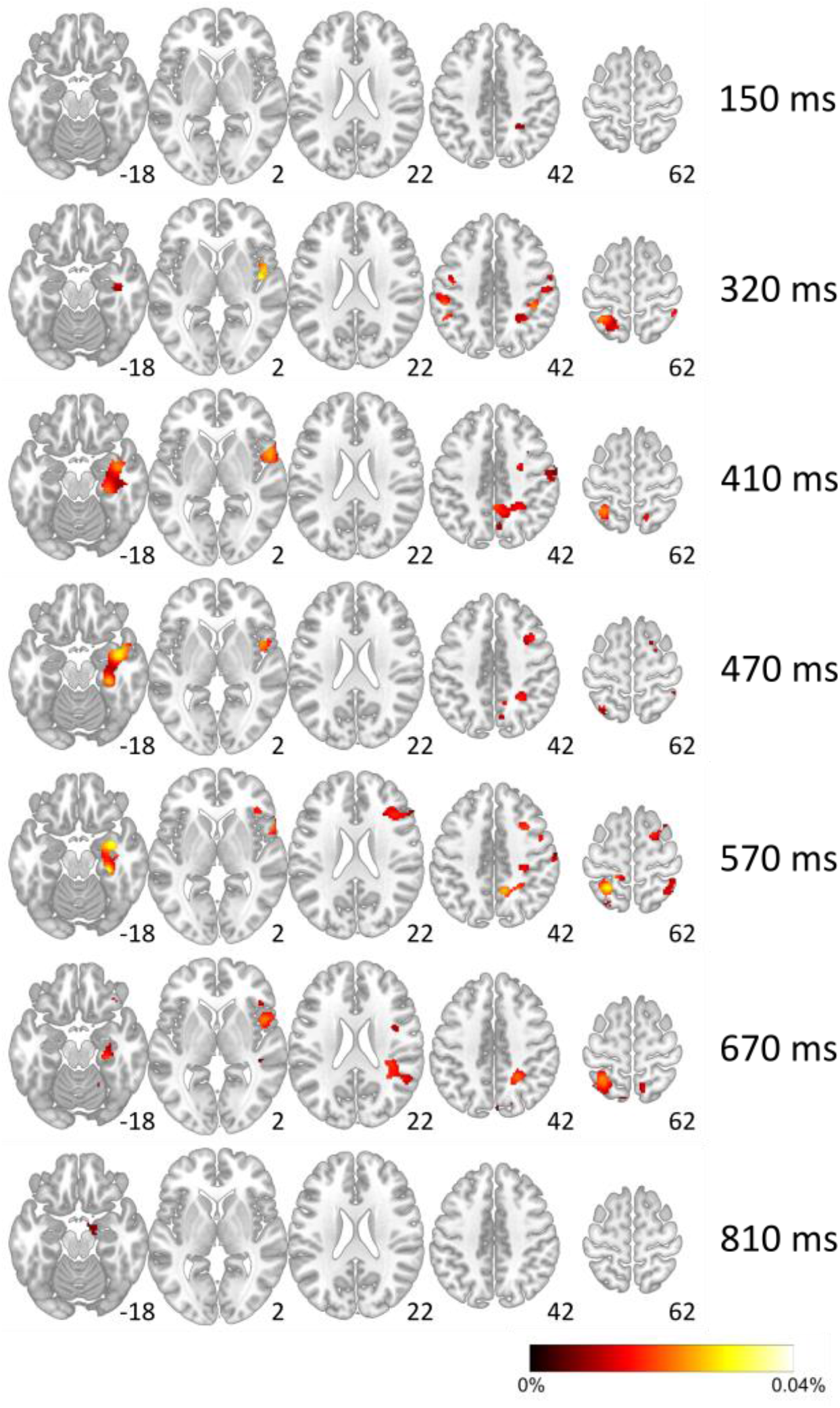
Axial slices exemplifying the results of the identity-model-based EEG-fMRI fusion analysis without a spatial and temporal extent threshold at selected timepoints. Color scale indicates how much of the variance of the correlation between the EEG RDM at this time point and the fMRI RDM at this voxel was explained by the identity model RDM (in percent) when controlling for pixel dissimilarity. A detailed animation of the full original spatio-temporal time course is available under www.cogsci.uni-jena.de/fmrieegfusion/. A customised online viewer is also included in the OSF repository.

B) Table of all spatial clusters within the significant spatio-temporal identity information time course

This table contains all spatial clusters within the significant spatio-temporal identity information time course after applying the spatial and temporal extent thresholds. It contains the MNI coordinates of the maximum voxel value in each spatial cluster and the anatomic labels that were automatically assigned to these coordinates by the SPM Anatomy toolbox as well as our manually revised labels. For the sake of transparency, note that the significant spatio-temporal time course also occasionally extended into the cerebellum as well as the brainstem. However, since we did not have sufficient fMRI-coverage of these regions, we decided not to interpret these results and excluded these clusters from the table.

Since our revised labels in this table are abbreviated, we hereby provide an overview of all abbreviations used.

Amy Amygdala

FusG Fusiform Gyrus

IFG Inferior Frontal Gyrus

Ins Insula

IPL Inferior Parietal Lobule

MFG Middle Frontal Gyrus

MTL Medial Temporal Lobe

PosG Postcentral Gyrus

Prec Precuneus

PreG Precentral Gyrus

SFG Superior Frontal Gyrus

SPL Superior Parietal Lobule

STG Superior Temporal Gyrus

TPJ Temporo-parietal Junction

**Table S2.**
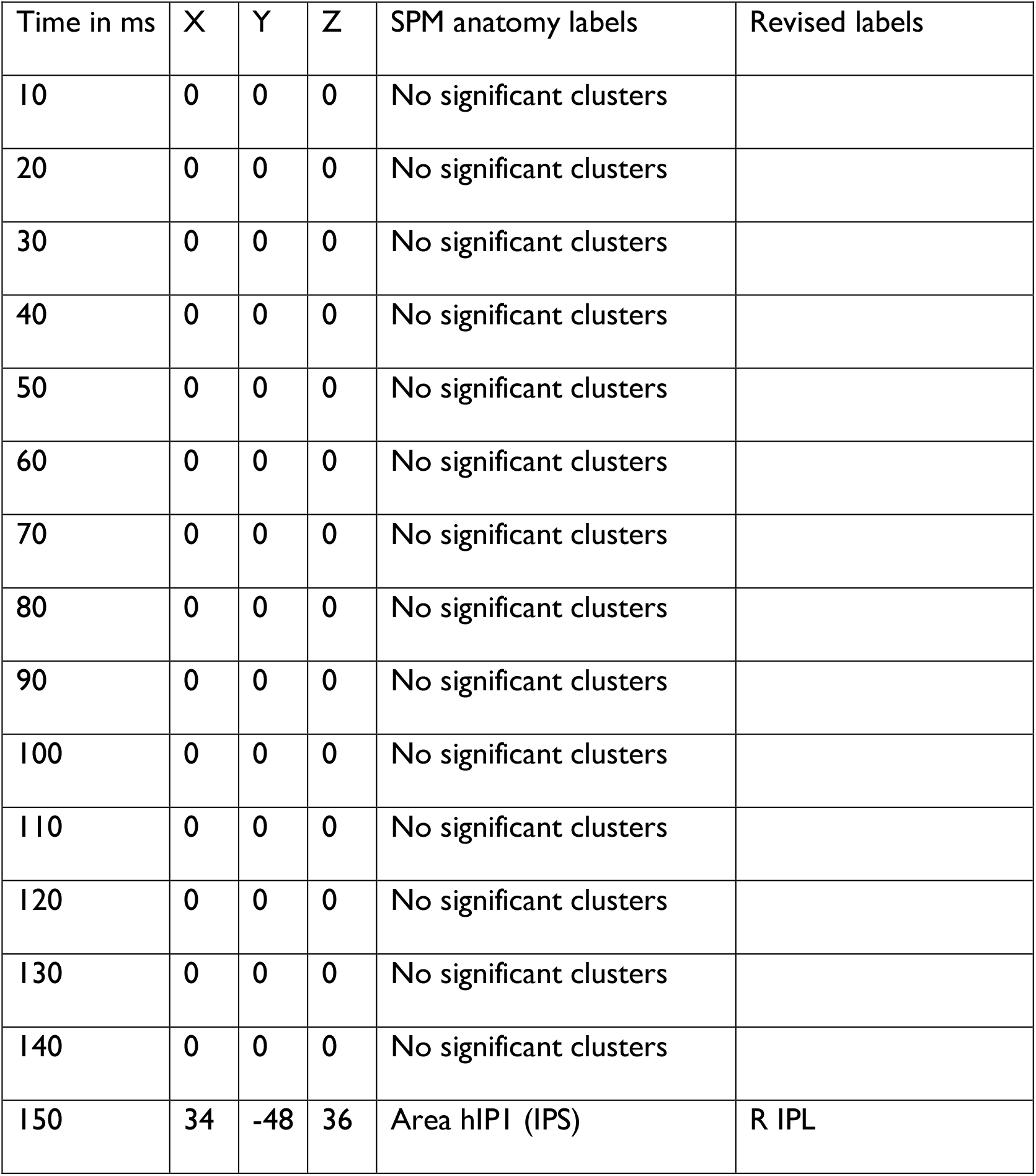
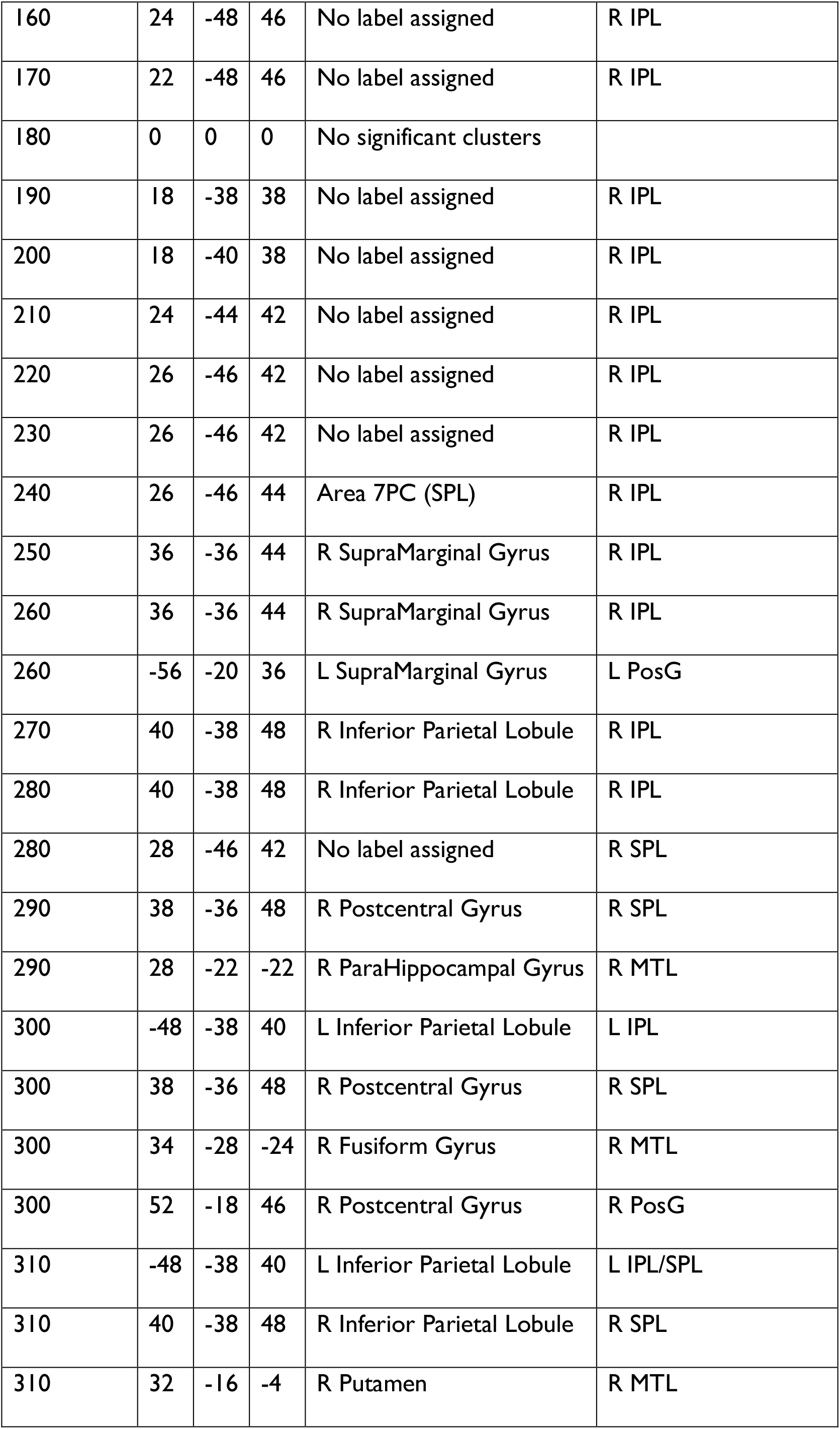
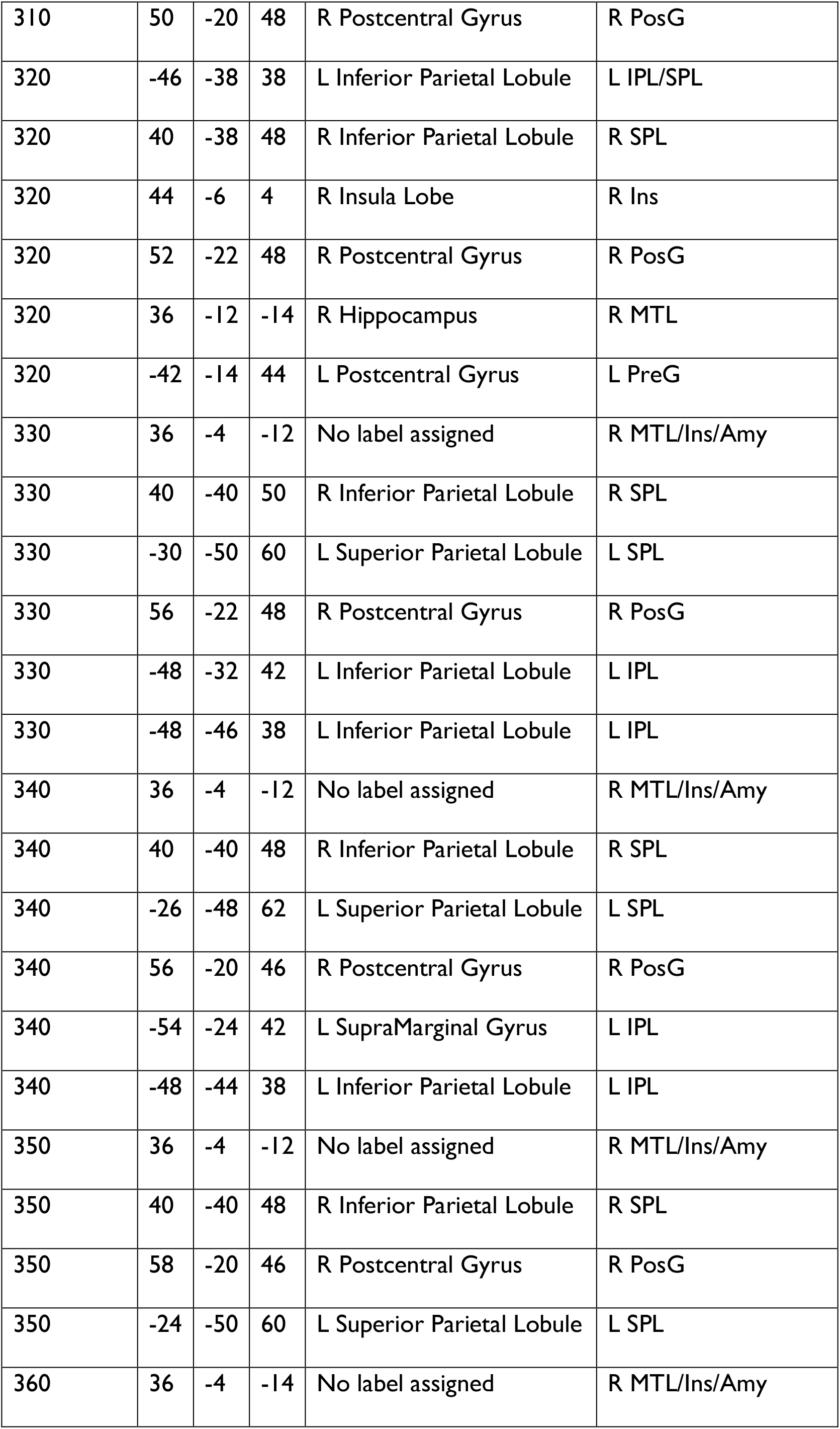
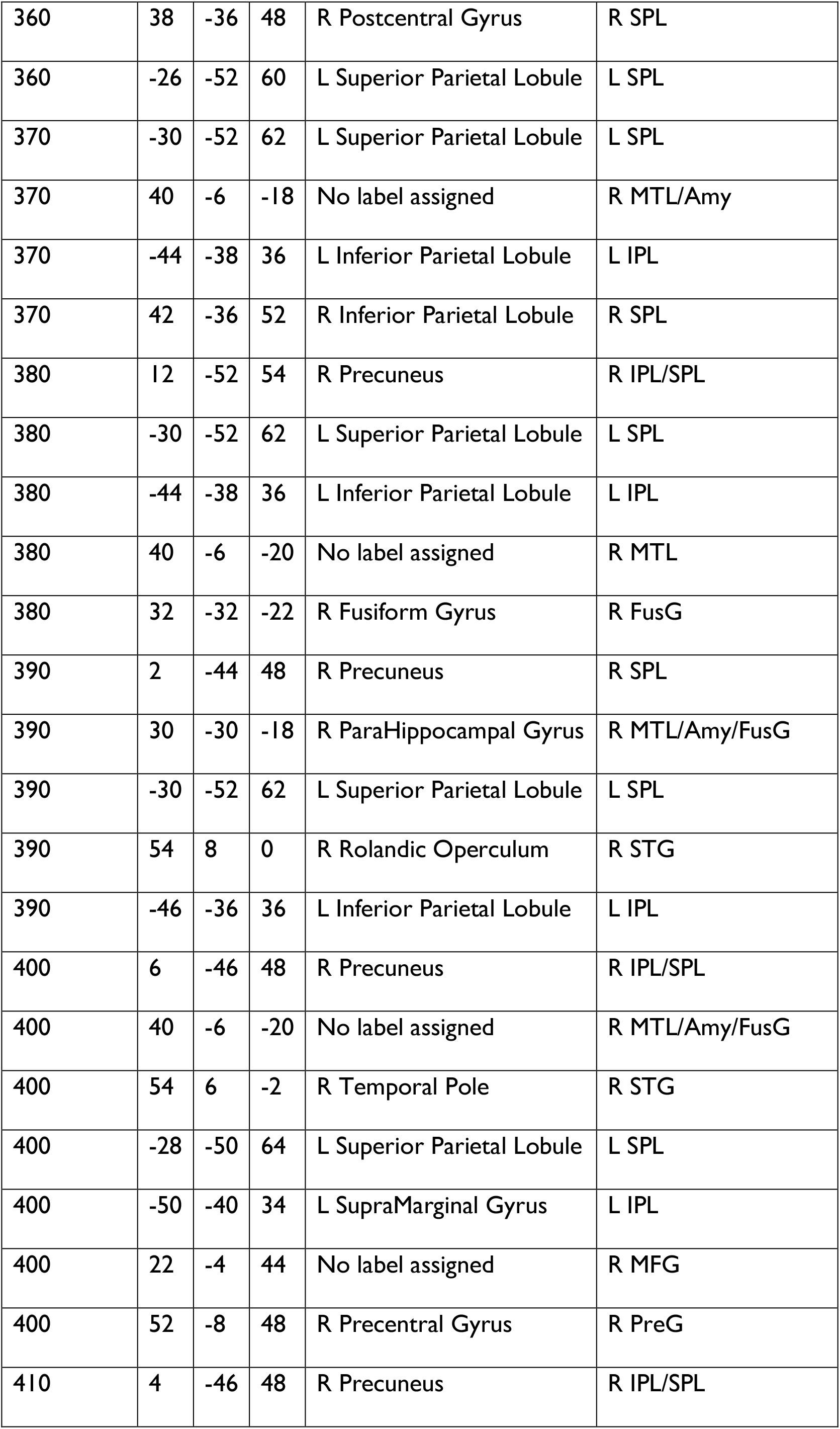
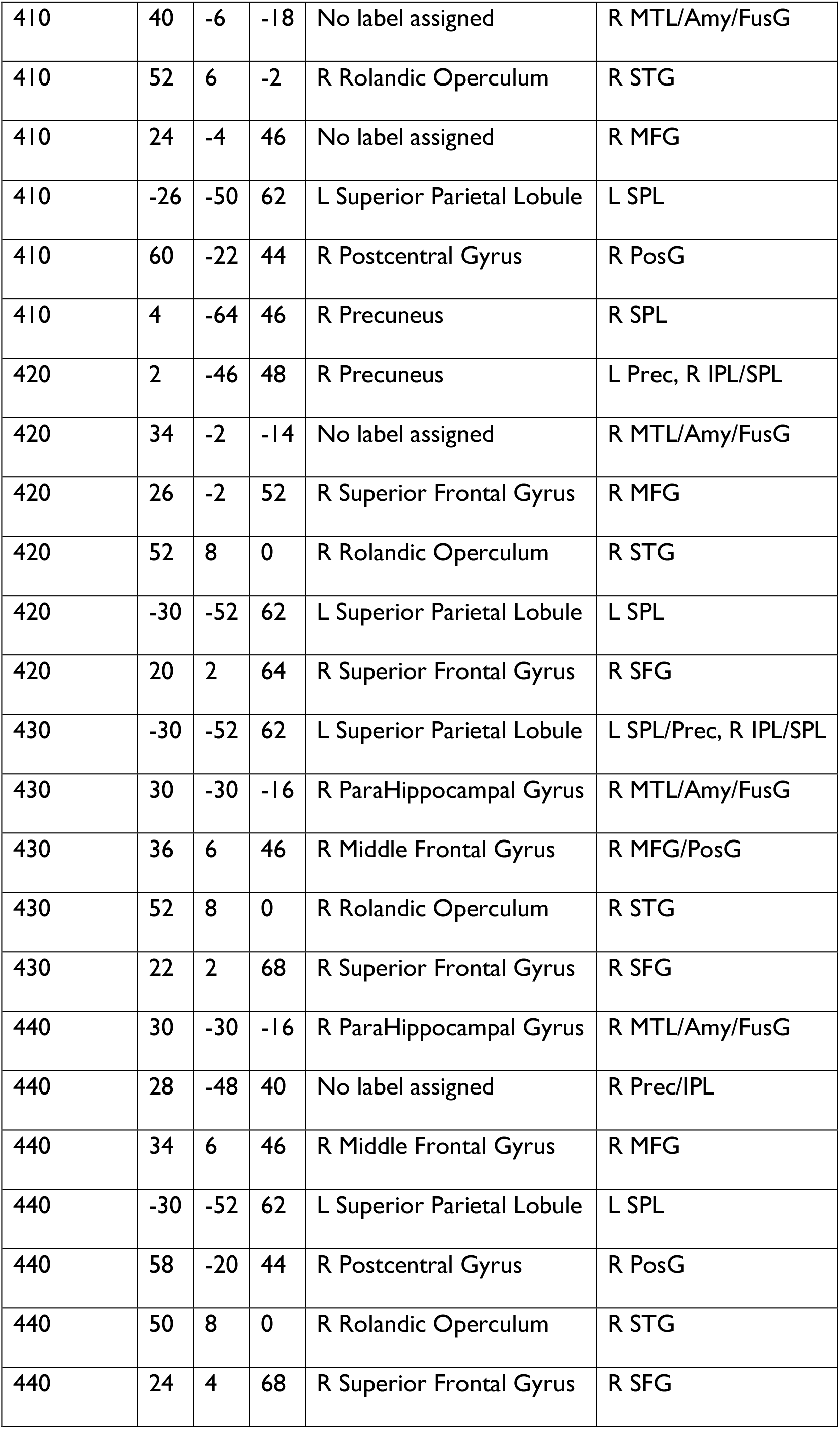
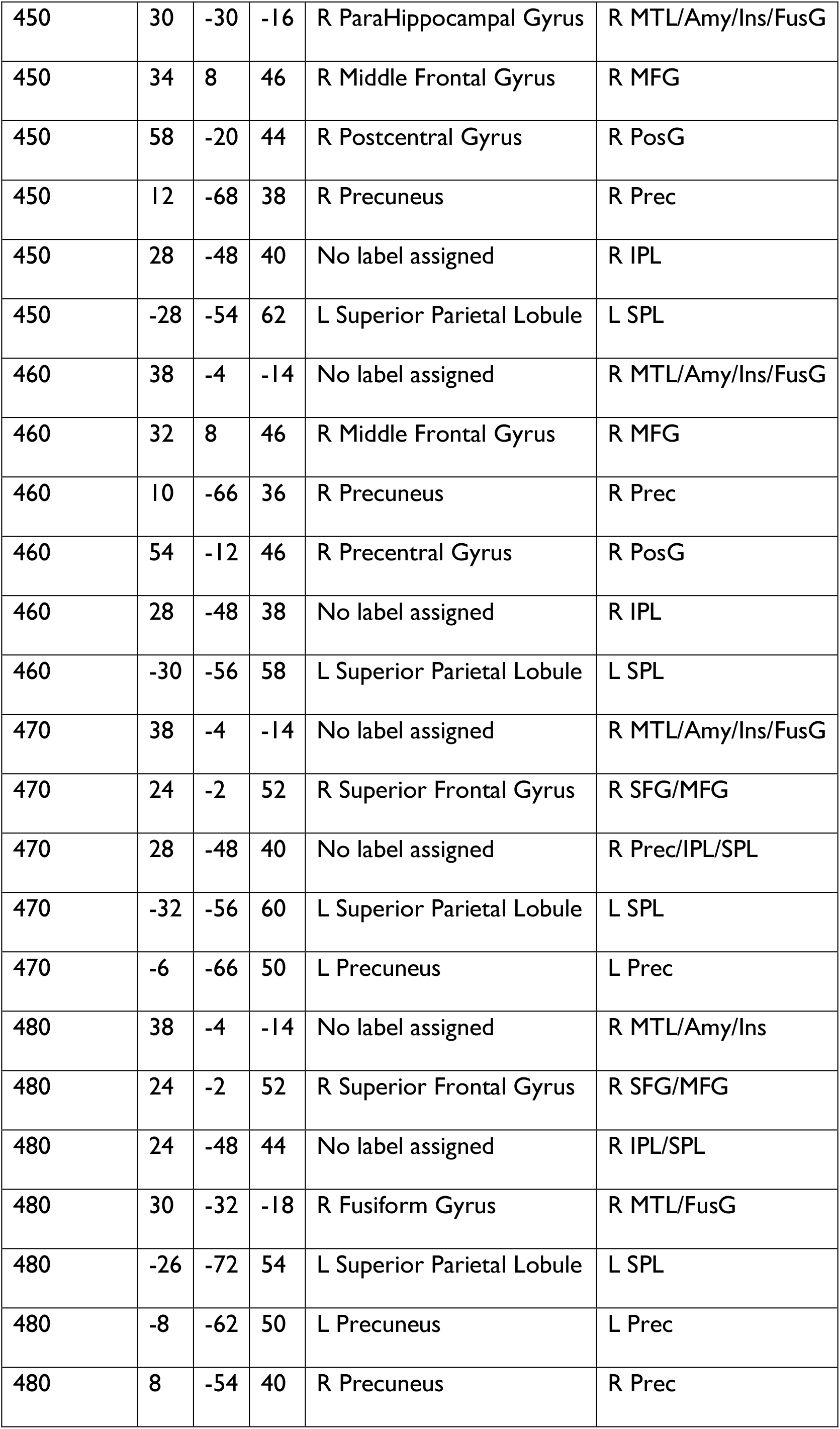
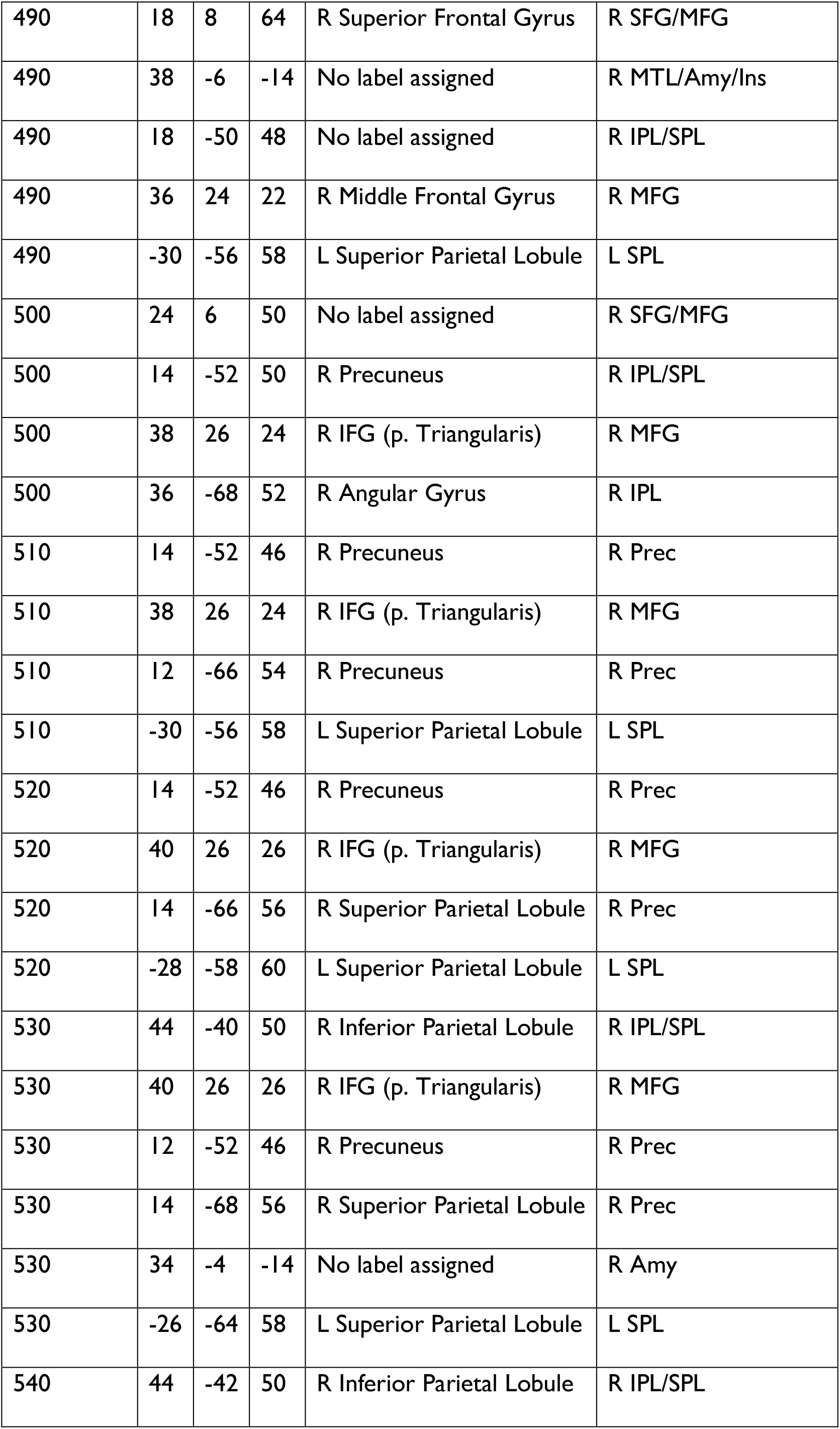
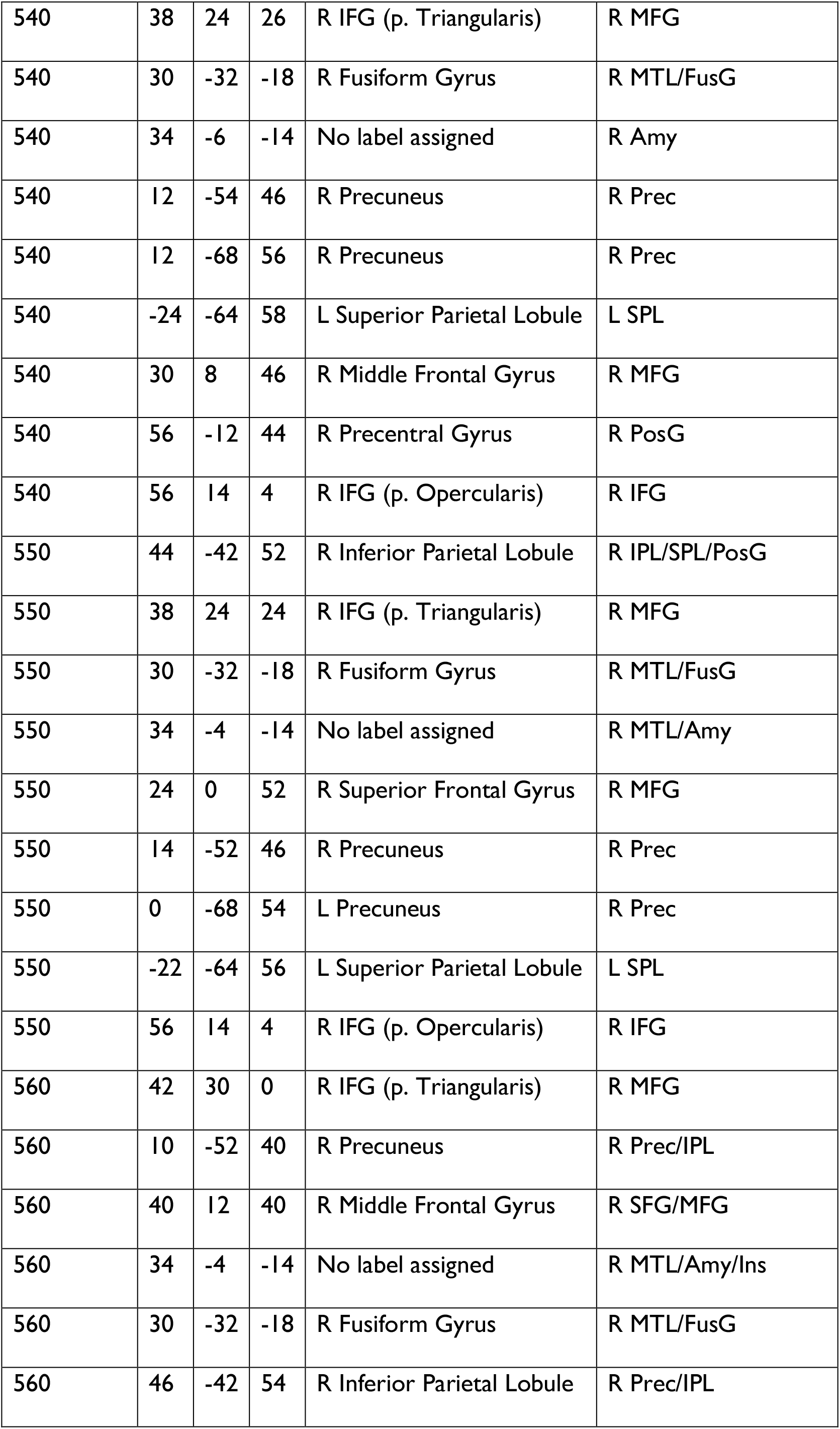
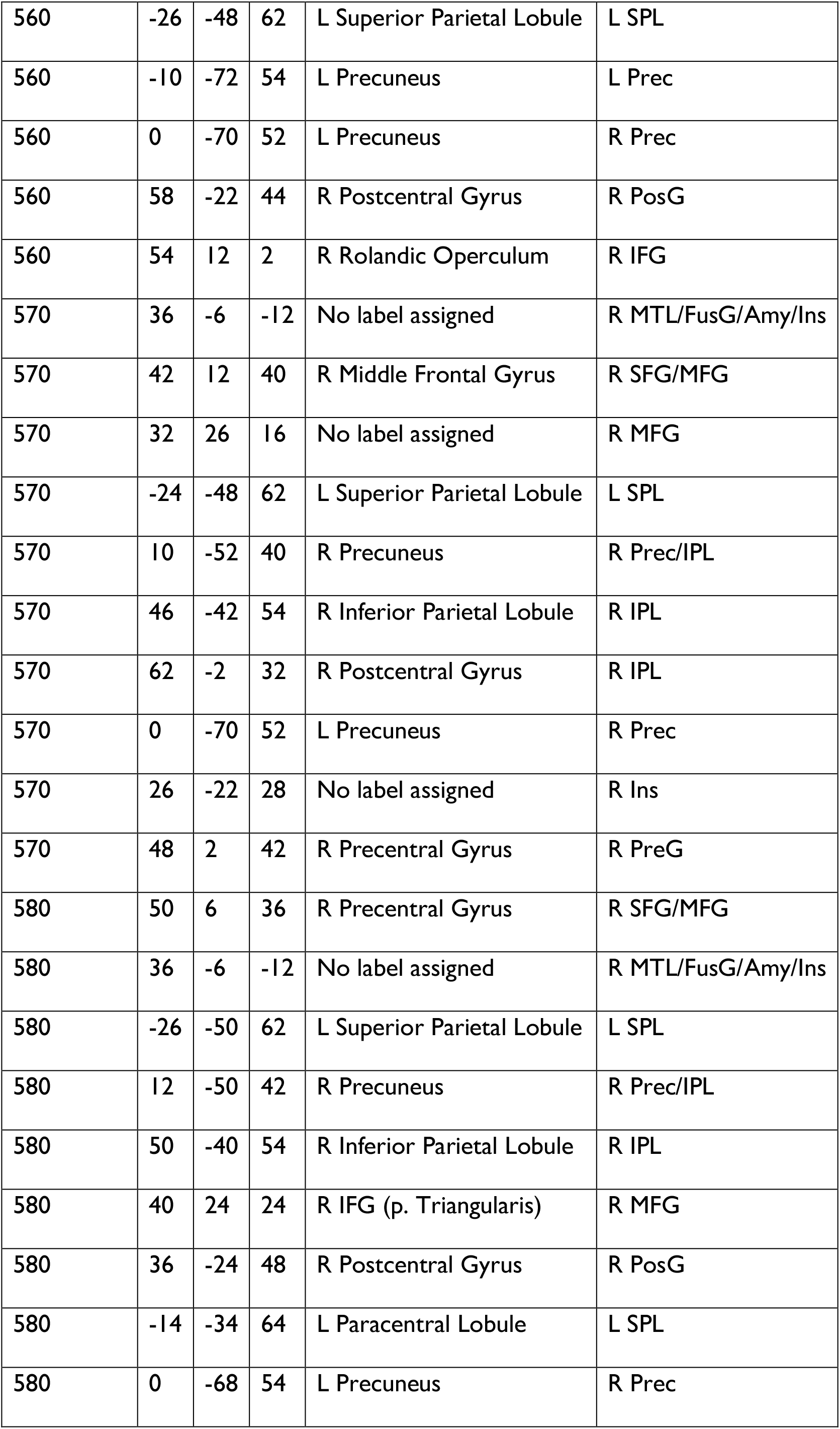
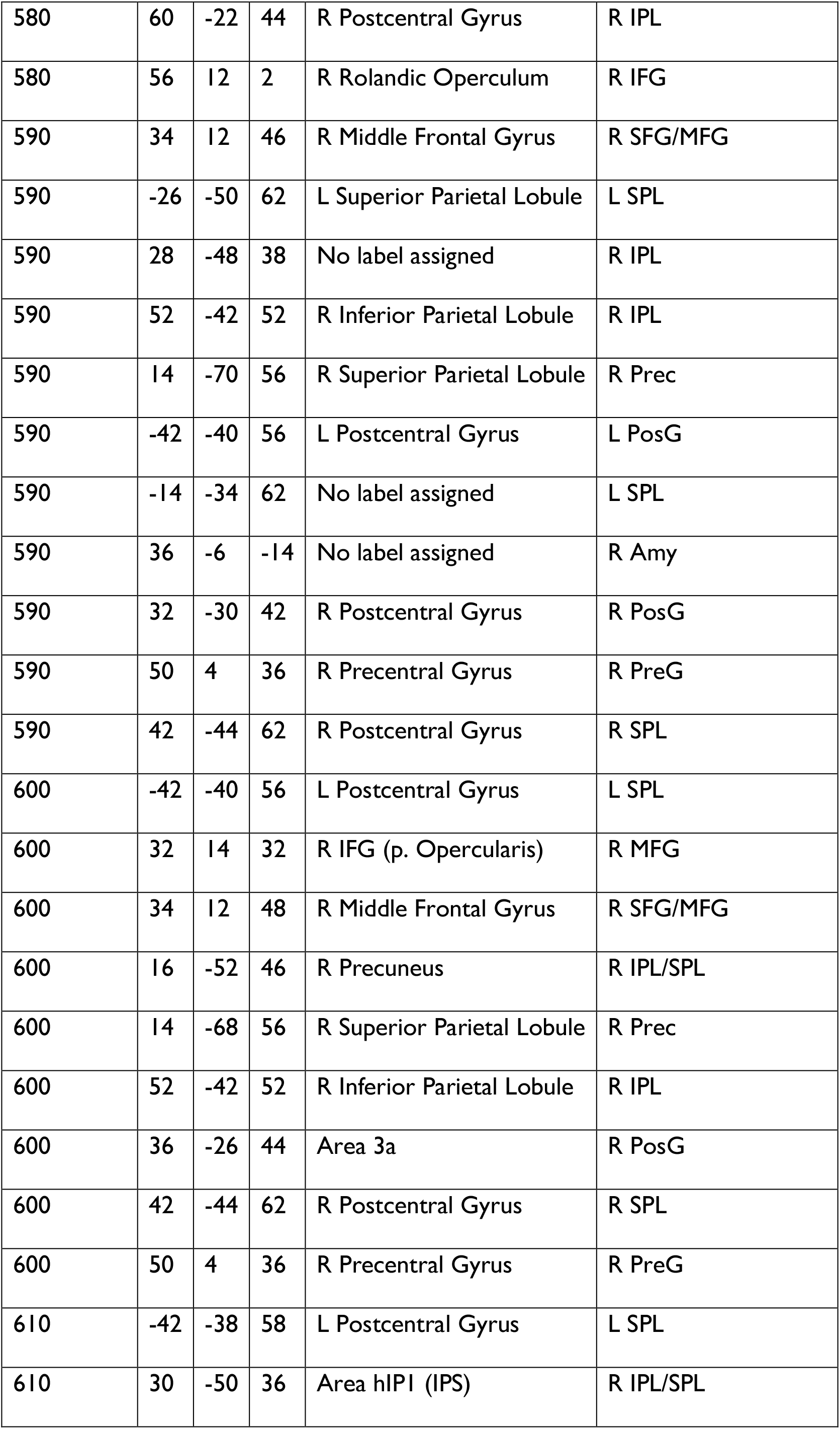
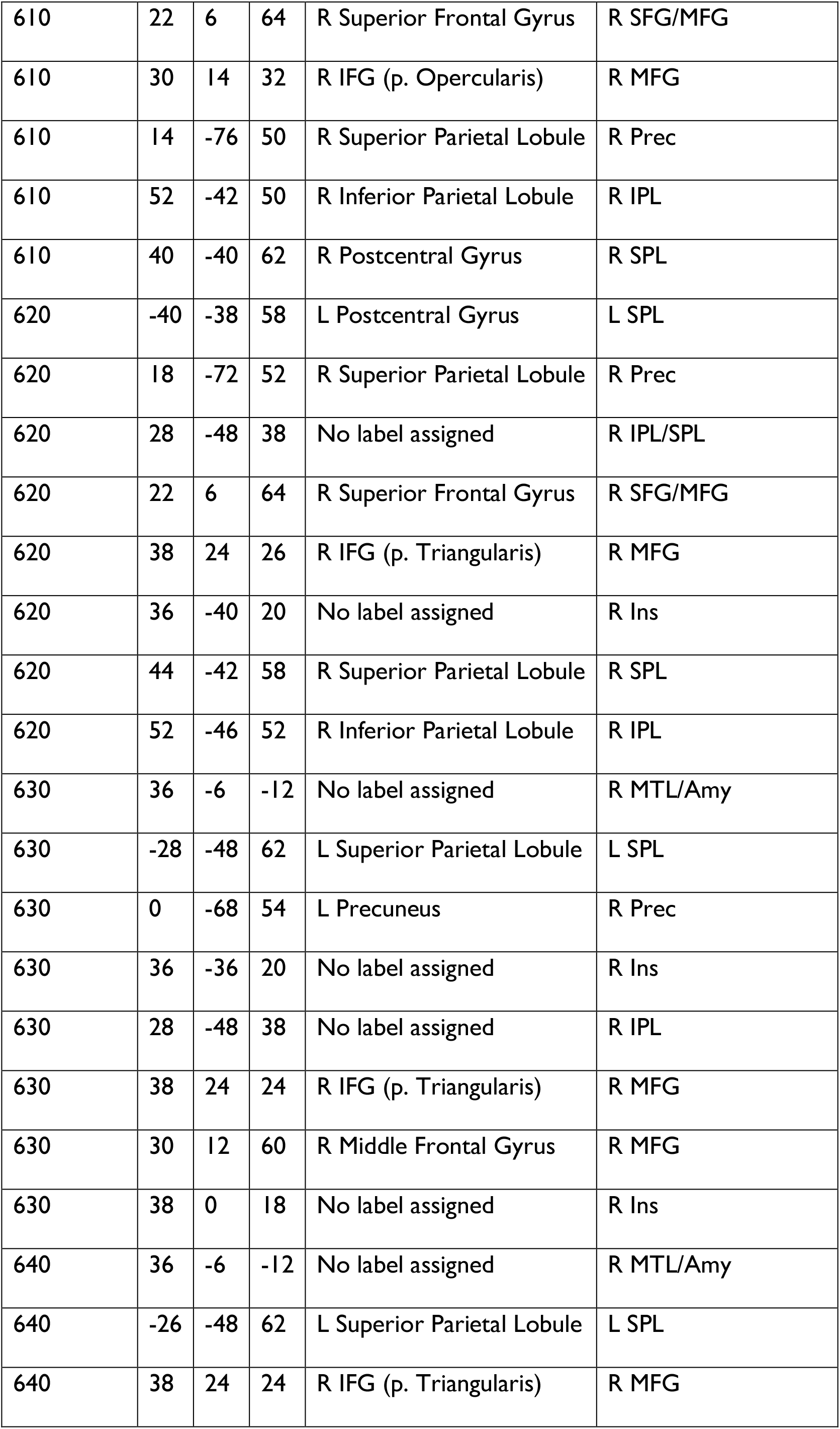
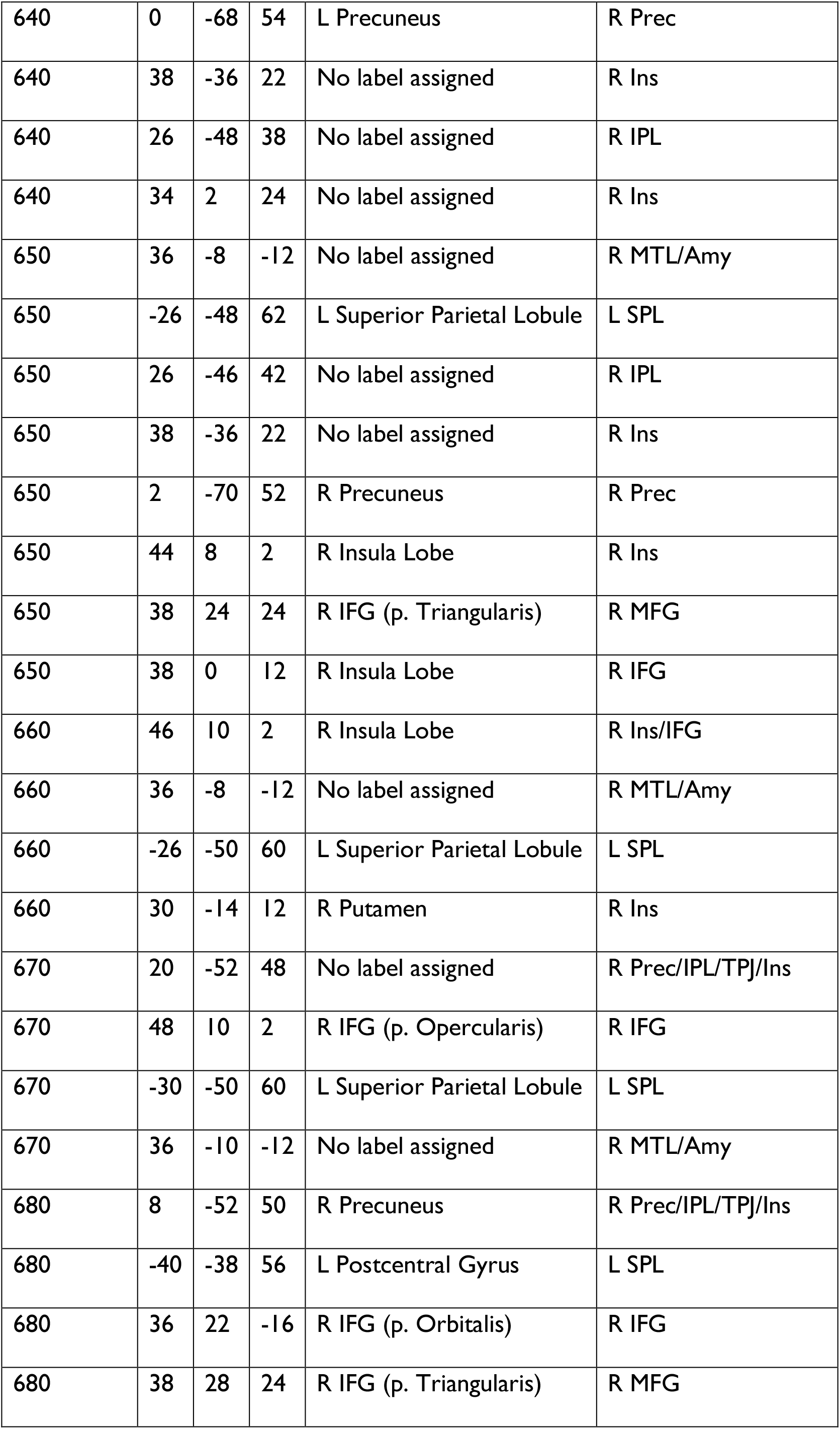
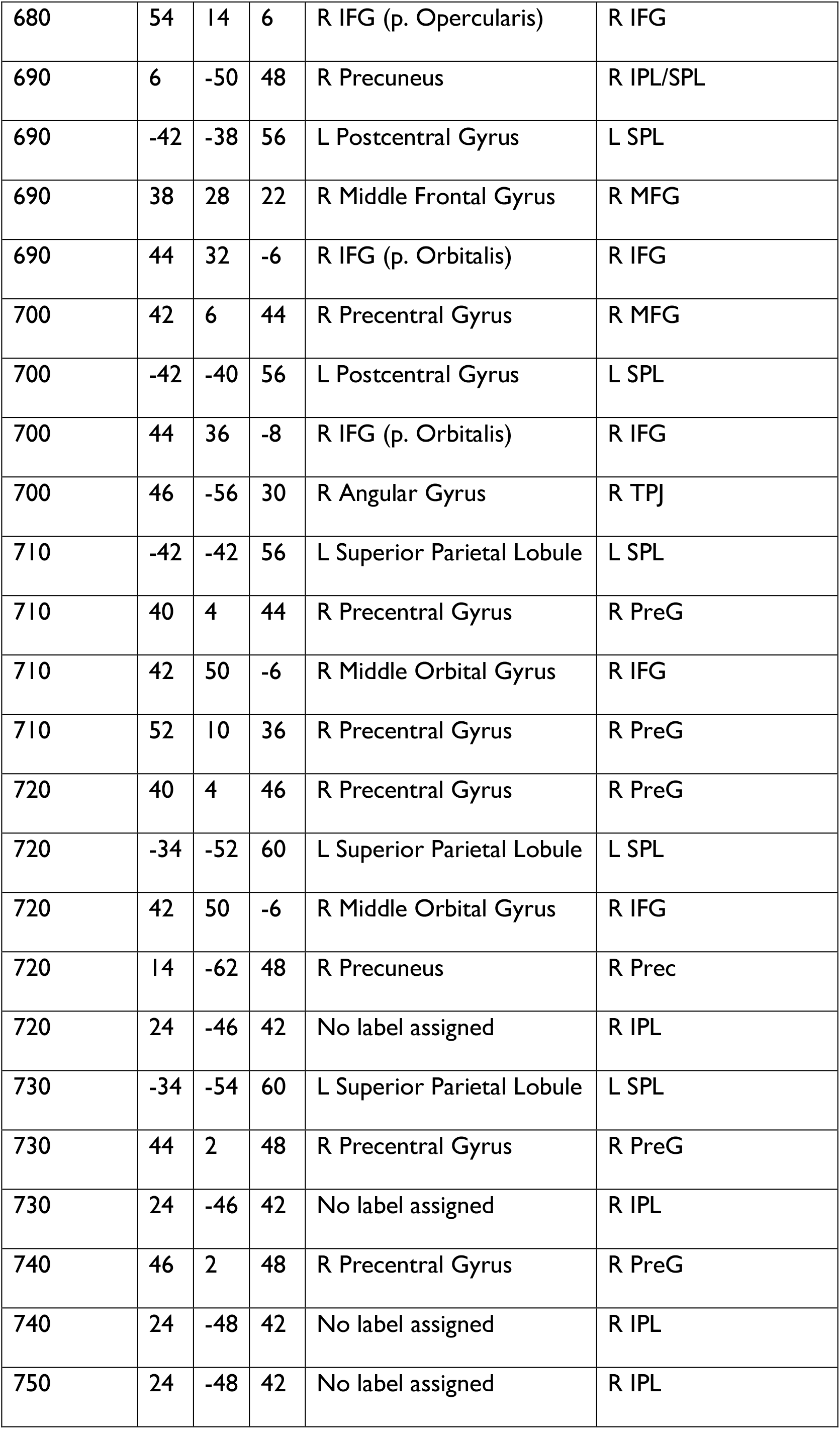
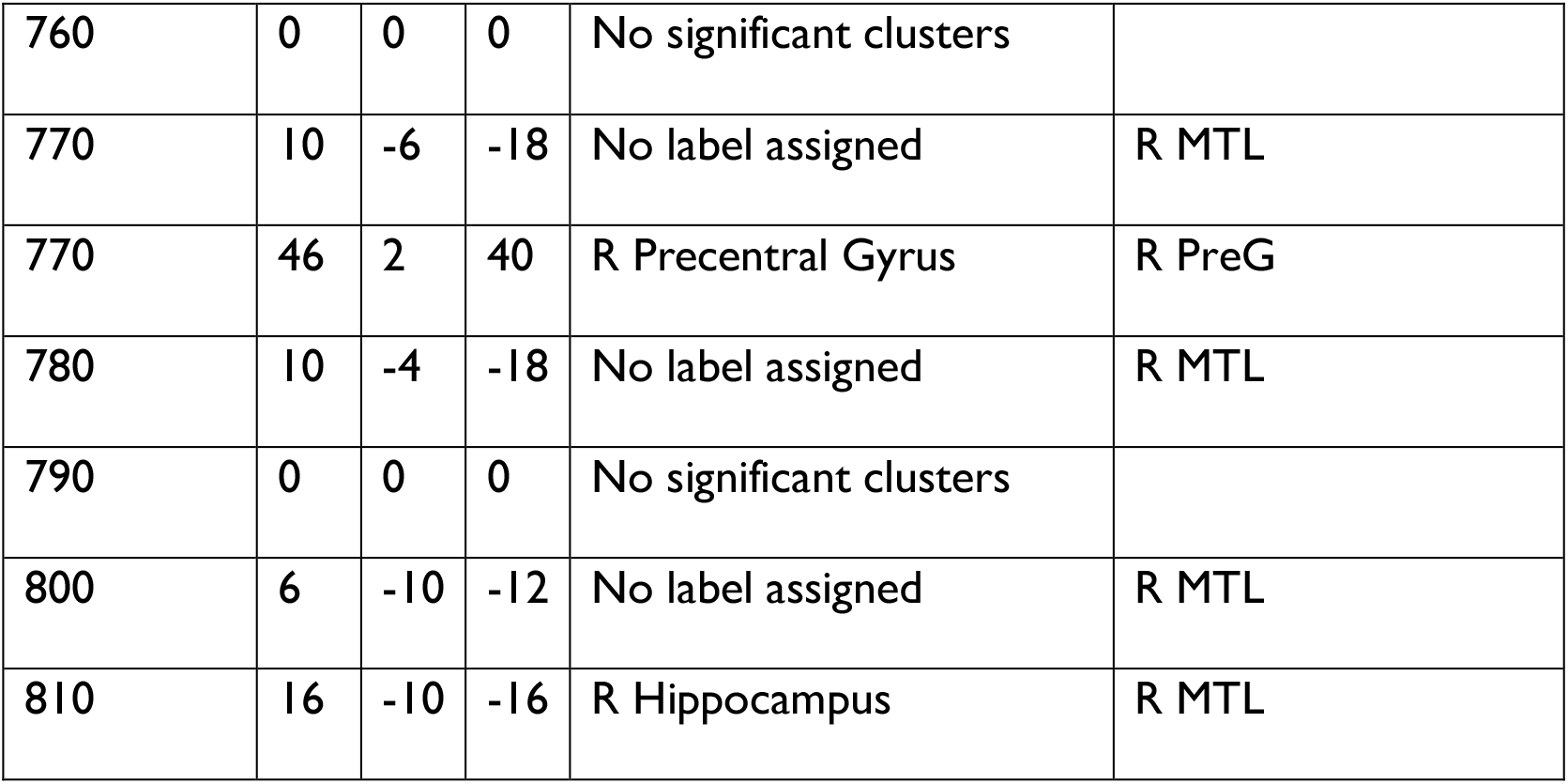
All spatial clusters within the significant spatio-temporal identity information time course with their MNI coordinates, SPM Anatomy toolbox labels and our revised versions of these labels.

C) Visualization of the significant spatio-temporal identity information time course

**Figure S3.**
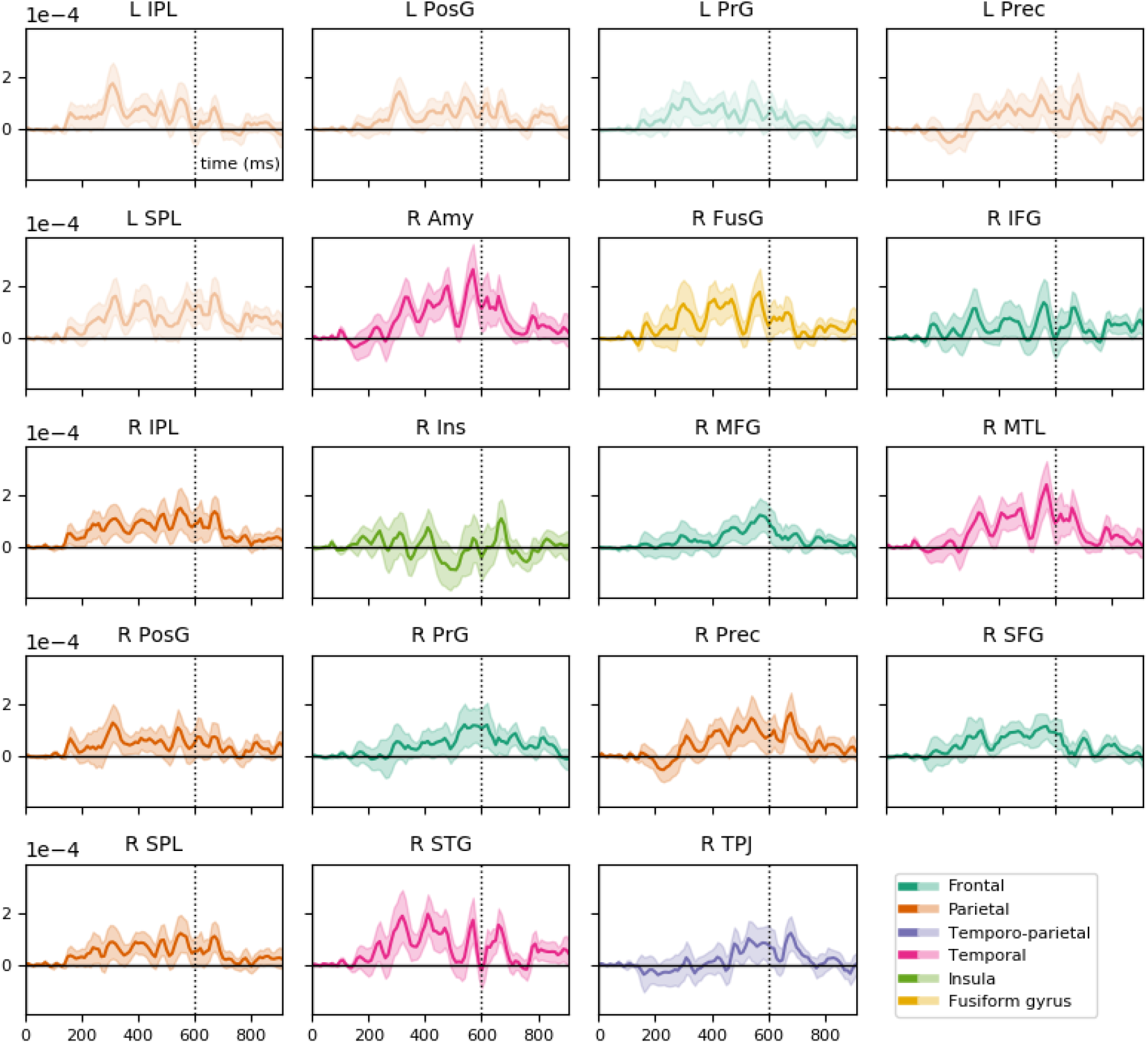
Mean identity commonality coefficients across time (with standard errors) from spheres (10 mm radius) inside each of the 19 regions in the significant spatio-temporal time course. For an animation of the full significant spatio-temporal time course with the spatial and temporal extent thresholds applied, see www.cogsci.uni-jena.de/fmrieegfusion/.

## References

Ambrus, G.G., Eick, C.M., Kaiser, D., Kovács, G., 2021. Getting to know you: emerging neural representations during face familiarization. J. Neurosci. JN-RM-2466–20. https://doi.org/10.1523/jneurosci.2466-20.2021

Ambrus, G.G., Kaiser, D., Cichy, R.M., Kovács, G., 2019. The Neural Dynamics of Familiar Face Recognition. Cereb. Cortex 29, 4775–4784. https://doi.org/10.1093/cercor/bhz010

Anzellotti, S., Fairhall, S.L., Caramazza, A., 2014. Decoding representations of face identity that are tolerant to rotation. Cereb. Cortex 24, 1988–1995. https://doi.org/10.1093/cercor/bht046

Axelrod, V., Yovel, G., 2015. Successful decoding of famous faces in the fusiform face area. PLoS One 10, 19–25. https://doi.org/10.1371/journal.pone.0117126

Axelrod, V., Yovel, G., 2013. The challenge of localizing the anterior temporal face area: A possible solution. Neuroimage 81, 371–380. https://doi.org/10.1016/j.neuroimage.2013.05.015

Bayer, M., Berhe, O., Dziobek, I., Johnstone, T., 2021. Rapid Neural Representations of Personally Relevant Faces. Cereb. Cortex. https://doi.org/10.1093/cercor/bhab116

Blank, H., Wieland, N., von Kriegstein, K., 2014. Person recognition and the brain: Merging evidence from patients and healthy individuals. Neurosci. Biobehav. Rev. 47, 717–734. https://doi.org/10.1016/j.neubiorev.2014.10.022

Bode, S., Feuerriegel, D., Bennett, D., Alday, P.M., 2019. The Decision Decoding ToolBOX (DDTBOX) – A Multivariate Pattern Analysis Toolbox for Event-Related Potentials. Neuroinformatics 17, 27–42. https://doi.org/10.1007/s12021-018-9375-z

Brainard, D.H., 1997. The Psychophysics Toolbox. Spat. Vis. 10, 433–436. https://doi.org/10.1163/156856897X00357

Burgess, N., Maguire, E.A., Spiers, H.J., O’Keefe, J., 2001. A temporoparietal and prefrontal network for retrieving the spatial context of lifelike events. Neuroimage 14, 439–453. https://doi.org/10.1006/nimg.2001.0806

Chan, A.W.-Y., Downing, P.E., 2011. Faces and Eyes in Human Lateral Prefrontal Cortex. Front. Hum. Neurosci. 5, 51. https://doi.org/10.3389/fnhum.2011.00051

Chang, C.C., Lin, C.J., 2011. LIBSVM: A Library for support vector machines. ACM Trans. Intell. Syst. Technol. 2, 1–39. https://doi.org/10.1145/1961189.1961199

Chang, D.H.F., Troje, N.F., Ikegaya, Y., Fujita, I., Ban, H., 2021. Spatiotemporal dynamics of responses to biological motion in the human brain. Cortex 136, 124–139. https://doi.org/10.1016/j.cortex.2020.12.015

Cichy, R.M., Pantazis, D., 2017. Multivariate pattern analysis of MEG and EEG: A comparison of representational structure in time and space. Neuroimage 158, 441–454. https://doi.org/10.1016/j.neuroimage.2017.07.023

Cichy, R.M., Pantazis, D., Oliva, A., 2016. Similarity-Based Fusion of MEG and fMRI Reveals Spatio-Temporal Dynamics in Human Cortex During Visual Object Recognition. Cereb. Cortex 26, 3563–3579. https://doi.org/10.1093/cercor/bhw135

Cichy, R.M., Pantazis, D., Oliva, A., 2014. Resolving human object recognition in space and time. Nat. Neurosci. 17, 455–462. https://doi.org/10.1038/nn.3635

Collins, J.A., Koski, J.E., Olson, I.R., 2016. More than meets the eye: The merging of perceptual and conceptual knowledge in the anterior temporal face area. Front. Hum. Neurosci. 10. https://doi.org/10.3389/fnhum.2016.00189

Collins, J.A., Olson, I.R., 2014. Beyond the FFA: The role of the ventral anterior temporal lobes in face processing. Neuropsychologia 61, 65–79. https://doi.org/10.1016/j.neuropsychologia.2014.06.005

Delorme, A., Makeig, S., 2004. EEGLAB: An open source toolbox for analysis of single-trial EEG dynamics including independent component analysis. J. Neurosci. Methods 134, 9– 21. https://doi.org/10.1016/j.jneumeth.2003.10.009

Dima, D.C., Perry, G., Singh, K.D., 2018. Spatial frequency supports the emergence of categorical representations in visual cortex during natural scene perception. Neuroimage 179, 102–116. https://doi.org/10.1016/j.neuroimage.2018.06.033

Dobs, K., Isik, L., Pantazis, D., Kanwisher, N., 2019. How face perception unfolds over time. Nat. Commun. 10, 1258. https://doi.org/10.1038/s41467-019-09239-1

Duchaine, B., Yovel, G., 2015. A Revised Neural Framework for Face Processing. Annu. Rev. Vis. Sci. 1, 393–416. https://doi.org/10.1146/annurev-vision-082114-035518

Frith, C.D., Frith, U., 1999. Interacting Minds-A Biological Basis. Sci. Compass 286.

Gilaie-Dotan, S., Malach, R., 2007. Sub-exemplar shape tuning in human face-related areas. Cereb. Cortex 17, 325–338. https://doi.org/10.1093/cercor/bhj150

Gilmore, A.W., Nelson, S.M., McDermott, K.B., 2015. A parietal memory network revealed by multiple MRI methods. Trends Cogn. Sci. 19, 534–543. https://doi.org/10.1016/j.tics.2015.07.004

Giordano, B.L., Whiting, C., Kriegeskorte, N., Kotz, S.A., Gross, J., Belin, P., 2021. The representational dynamics of perceived voice emotions evolve from categories to dimensions. Nat. Hum. Behav. https://doi.org/10.1038/s41562-021-01073-0

Gobbini, M.I., Haxby, J. V., 2007. Neural systems for recognition of familiar faces. Neuropsychologia 45, 32–41. https://doi.org/10.1016/j.neuropsychologia.2006.04.015

Gobbini, M.I., Haxby, J. V., 2006. Neural response to the visual familiarity of faces. Brain Res. Bull. 71, 76–82. https://doi.org/10.1016/j.brainresbull.2006.08.003

Goesaert, E., Op de Beeck, H.P., 2013. Representations of facial identity information in the ventral visual stream investigated with multivoxel pattern analyses. J. Neurosci. 33, 8549–8558. https://doi.org/10.1523/JNEUROSCI.1829-12.2013

Gorno-Tempini, M.L., Price, C.J., 2001 Identification of famous faces and buildings: a functional neuroimaging study of semantically unique items. Brain. 124(Pt 10), 2087–97. doi: 10.1093/brain/124.10.2087. PMID: 11571224.

Grootswagers, T., Wardle, S.G., Carlson, T.A., 2017. Decoding dynamic brain patterns from evoked responses: A tutorial on multivariate pattern analysis applied to time series neuroimaging data. J. Cogn. Neurosci. 29, 677–697. https://doi.org/10.1162/jocn_a_01068

Guntupalli, J.S., Wheeler, K.G., Gobbini, M.I., 2017. Disentangling the Representation of Identity from Head View Along the Human Face Processing Pathway. Cereb. Cortex 27, 46–53. https://doi.org/10.1093/cercor/bhw344

Haxby, J. V., Hoffman, E.A., Gobbini, M.I., 2000. The distributed human neural system for face perception. Trends Cogn. Sci. 4, 223–233. https://doi.org/10.1016/S1364-6613(00)01482-0

Hebart, M.N., Bankson, B.B., Harel, A., Baker, C.I., Cichy, R.M., 2018. The representational dynamics of task and object processing in humans. Elife 7, 1–21. https://doi.org/10.7554/eLife.32816

Hebart, M.N., Görgen, K., Haynes, J.-D., Dubois, J., 2015. The Decoding Toolbox (TDT): a versatile software package for multivariate analyses of functional imaging data. Front. Neuroinform. 8, 1–18. https://doi.org/10.3389/fninf.2014.00088

Ida Gobbini, M., Leibenluft, E., Santiago, N., Haxby, J. V., 2004. Social and emotional attachment in the neural representation of faces. Neuroimage 22, 1628–1635. https://doi.org/10.1016/j.neuroimage.2004.03.049

Jenkins, R., White, D., Van Montfort, X., Mike Burton, A., 2011. Variability in photos of the same face. Cognition 121, 313–323. https://doi.org/10.1016/j.cognition.2011.08.001

Jeong, S.K., Xu, Y., 2016. Behaviorally Relevant Abstract Object Identity Representation in the Human Parietal Cortex. https://doi.org/10.1523/JNEUROSCI.1016-15.2016

Kleiner, M., Brainard, D., Pelli, D., Ingling, A., Murray, R., Broussard, C., 2007. What’s new in psychtoolbox-3, Perception. [Pion Ltd.].

Kovács, G., 2020. Getting to know someone: Familiarity, person recognition, and identification in the human brain. J. Cogn. Neurosci. 32, 2205–2225. https://doi.org/10.1162/jocn_a_01627

Kriegeskorte, N., Goebel, R., Bandettini, P., 2006. Information-based functional brain mapping. Proc. Natl. Acad. Sci. U. S. A. 103, 3863–3868.

Kriegeskorte, N., Formisano, E., Sorger, B., Goebel, R., 2007. Individual faces elicit distinct response patterns in human anterior temporal cortex. Proc. Natl. Acad. Sci. U. S. A. 104, 20600–20605. https://doi.org/10.1073/pnas.0705654104

Kriegeskorte, N., Kievit, R.A., 2013. Representational geometry: Integrating cognition, computation, and the brain. Trends Cogn. Sci. 17, 401–412. https://doi.org/10.1016/j.tics.2013.06.007

Leech, R., Sharp, D.J., 2014. The role of the posterior cingulate cortex in cognition and disease Abbreviations: ADHD = attention deficit hyperactivity disorder; DMN = default mode network; FPCN = fronto-parietal control network; PCC = posterior cingulate cortex. A J. Neurol. https://doi.org/10.1093/brain/awt162

Morton, N.W., Zippi, E.L., Noh, S.M., Preston, A.R., 2021. Semantic knowledge of famous people and places is represented in hippocampus and distinct cortical networks. J. Neurosci. 41, 2762–2779. https://doi.org/10.1523/JNEUROSCI.2034-19.2021

Mullen, T., 2012. CleanLine EEGLAB plugin. San Diego, CA Neuroimaging Informatics Toolsand Resour. Clear.

Muukkonen, I., Ölander, K., Numminen, J., Salmela, V.R., 2020. Spatio-temporal dynamics of face perception. Neuroimage 209, 116531. https://doi.org/10.1016/j.neuroimage.2020.116531

Muukkonen, I., Ölander, K., Salmela, V.R., 2019. Spatio-temporal dynamics of face perception. bioRxiv 1–23. https://doi.org/10.1101/550038

Nasr, S., Tootell, R.B.H., 2012. Role of fusiform and anterior temporal cortical areas in facial recognition. Neuroimage 63, 1743–1753. https://doi.org/10.1016/j.neuroimage.2012.08.031

Natu, V., O’Toole, A.J., 2011. The neural processing of familiar and unfamiliar faces: A review and synopsis. Br. J. Psychol. 102, 726–747. https://doi.org/10.1111/j.2044-8295.2011.02053.x

Nemrodov, D., Niemeier, M., Mok, J.N.Y., Nestor, A., 2016. The time course of individual face recognition: A pattern analysis of ERP signals. Neuroimage 132, 469–476. https://doi.org/10.1016/j.neuroimage.2016.03.006

Nemrodov, D., Niemeier, M., Patel, A., Nestor, A., 2018. The neural dynamics of facial identity processing: Insights from EEG-based pattern analysis and image reconstruction. eNeuro 5. https://doi.org/10.1523/ENEURO.0358-17.2018

Nestor, A., Plaut, D.C., Behrmann, M., 2011. Unraveling the distributed neural code of facial identity through spatiotemporal pattern analysis. Proc. Natl. Acad. Sci. 108, 9998– 10003. https://doi.org/10.1073/pnas.1102433108

Nichols, T.E., Holmes, A.P., 2002. Nonparametric permutation tests for functional neuroimaging: A primer with examples. Hum. Brain Mapp. 15, 1–25. https://doi.org/10.1002/hbm.1058

Peirce, J.W., 2008. Generating stimuli for neuroscience using PsychoPy. Front. Neuroinform.

Pelli, D.G., 1997. The VideoToolbox software for visual psychophysics: transforming numbers into movies. Spat. Vis. 10, 437–42.

Rajimehr, R., Young, J.C., Tootell, R.B.H., 2009. An anterior temporal face patch in human cortex, predicted by macaque maps. Proc. Natl. Acad. Sci. 106, 1995–2000. https://doi.org/10.1073/pnas.0807304106

Ramon, M., Vizioli, L., Liu-Shuang, J., Rossion, B., 2015. Neural microgenesis of personally familiar face recognition. Proc. Natl. Acad. Sci. 112. https://doi.org/10.1073/pnas.1414929112

Ritchie, J.B., Bracci, S., Op de Beeck, H., 2017. Avoiding illusory effects in representational similarity analysis: What (not) to do with the diagonal. Neuroimage. https://doi.org/10.1016/j.neuroimage.2016.12.079

Ruxton, G.D., Neuhäuser, M., 2013. Improving the reporting of P-values generated by randomization methods. Methods Ecol. Evol. 4, 1033–1036. https://doi.org/10.1111/2041-210X.12102

Seibold, D.R., McPhee, R.D., 1979. Commonality Analysis: a Method for Decomposing Explained Variance in Multiple Regression Analyses. Hum. Commun. Res. 5, 355–365. https://doi.org/10.1111/j.1468-2958.1979.tb00649.x

Tsantani, M., Kriegeskorte, N., McGettigan, C., Garrido, L., 2019. Faces and voices in the brain: A modality-general person-identity representation in superior temporal sulcus. Neuroimage 201, 116004. https://doi.org/10.1016/j.neuroimage.2019.07.017

Tsantani, M., Kriegeskorte, N., Storrs, K., Williams, A.L., McGettigan, C., Garrido, L., 2021. FFA and OFA encode distinct types of face identity information. J. Neurosci. 41, JN-RM-1449-20. https://doi.org/10.1523/jneurosci.1449-20.2020

Verosky, S.C., Todorov, A., Turk-Browne, N.B., 2013. Representations of individuals in ventral temporal cortex defined by faces and biographies. Neuropsychologia 51, 2100– 2108. https://doi.org/10.1016/j.neuropsychologia.2013.07.006

Vida, M.D., Nestor, A., Plaut, D.C., Behrmann, M., 2017. Spatiotemporal dynamics of similarity-based neural representations of facial identity. Proc. Natl. Acad. Sci. U. S. A. 114, 388–393. https://doi.org/10.1073/pnas.1614763114

Visconti Di Oleggio Castello, M., Halchenko, Y.O., Guntupalli, J.S., Gors, J.D., Gobbini, M.I., 2017. The neural representation of personally familiar and unfamiliar faces in the distributed system for face perception. Sci. Rep. 7, 1–14. https://doi.org/10.1038/s41598-017-12559-1

Wagner, A.D., Shannon, B.J., Kahn, I., Buckner, R.L., 2005. Parietal lobe contributions to episodic memory retrieval. Trends Cogn. Sci. 9, 445–453. https://doi.org/10.1016/j.tics.2005.07.001

Watson, R., Huis In’t Veld, E.M.J., de Gelder, B., 2016. The neural basis of individual face and object perception. Front. Hum. Neurosci. 10. https://doi.org/10.3389/fnhum.2016.00066

Weibert, K., Harris, R.J., Mitchell, A., Byrne, H., Young, A.W., Andrews, T.J., 2016. An image-invariant neural response to familiar faces in the human medial temporal lobe. Cortex 84, 34–42. https://doi.org/10.1016/j.cortex.2016.08.014

Wiese, H., Tüttenberg, S.C., Ingram, B.T., Chan, C.Y.X., Gurbuz, Z., Burton, A.M., Young, A.W., 2019. A Robust Neural Index of High Face Familiarity. Psychol. Sci. 30, 261– 272. https://doi.org/10.1177/0956797618813572

Xu, Y., Chun, M.M., 2006. Dissociable neural mechanisms supporting visual short-term memory for objects. Nature 440, 91–95. https://doi.org/10.1038/nature04262

